# Mechanism of GTPase activation of a prokaryotic small Ras-like GTPase MglA by an asymmetrically interacting MglB dimer

**DOI:** 10.1101/2023.03.05.531159

**Authors:** Sukanya Chakraborty, Manil Kanade, Pananghat Gayathri

**Affiliations:** Indian Institute of Science Education and Research Pune India; IISER PUNE; Indian Institute of Science Education and Research Pune

**Keywords:** Prokaryotic small Ras-like GTPase MglA, MglB, asymmetry, GAP, Roadblock domain, GEF, allostery

## Abstract

Cell polarity oscillations in *Myxococcus xanthus* motility are driven by a prokaryotic small Ras-like GTPase, MglA, which switches from one cell pole to the other in response to extracellular signals. MglA dynamics is regulated by MglB, which functions both as a GAP (GTPase activating protein) and a GEF (guanine nucleotide exchange factor) for MglA. With an aim to dissect the role of asymmetry in the dual GAP and GEF activities of MglB, we generated a functional MglAB complex by co-expressing MglB with a linked construct of MglA and MglB. This strategy enabled us to generate mutations of individual MglB protomers (MglB_1_ linked to MglA or MglB_2_) and delineate their role in GEF and GAP activities. We establish that the C-terminal helix of MglB_1_, but not MglB_2_, stimulates nucleotide exchange through a site away from the nucleotide-binding pocket, confirming an allosteric mechanism. Interaction between the N-terminal β-strand of MglB_1_ and β_0_ of MglA is essential for the GEF activity of MglB. Specific residues of MglB_2_ which interact with Switch-I of MglA partially contribute to its GAP activity. Thus, the role of the MglB_2_ protomer in the GAP activity of MglB is limited to restricting the conformation of MglA active site loops by steric hindrance. The direct demonstration of the allosteric mechanism of GEF action provides us new insights into the regulation of small Ras-like GTPases, a feature potentially present in many uncharacterized GEFs.

## Introduction

Polarity determination and its regulation are critical to several vital cellular processes like signal transduction, cell growth and motility^1^. *Myxococcus xanthus* is a Gram-negative soil bacterium which has been used as a model organism to study polarity reversals that characterize its gliding motility pattern^2^. Polarity reversals are associated with spatial oscillations of the intracellular proteins as the bacterium switches its direction of movement. Frz chemosensory system and the small Ras-like GTPase MglA are two main players in the regulation of these oscillations^3–5^.

Small Ras-like GTPases play a key role in determining cell polarity^6^. These GTPases usually act as molecular switches alternating between its GTP-bound active conformation (‘ON’ state) and the GDP-bound inactive conformation (‘OFF’ state). This conformational transition reorients residues of the GTPase catalytic pocket, which constitutes the conserved Switch loops (Switch-I and Switch-II) which further signal downstream effectors. Most of these GTPases are associated with their respective GTPase Activating Proteins (GAPs), which assist GTP hydrolysis, and Guanine nucleotide Exchange Factors (GEFs), which stimulate nucleotide exchange^7^. MglA, a prototypic member of the prokaryotic small Ras-like GTPase family, localizes to the leading pole of the cell in its GTP-bound state, and the cell reverses polarity when it relocates^8,9^. MglB, which consists of a Roadblock/LC7 (Rbl) domain, acts as a GAP for MglA ^10^, and predominantly localizes to the opposite pole. We recently demonstrated that MglB also exerts a nucleotide exchange effect on MglA, thus performing the dual role of a GAP and a GEF ^11^. Such a dual mechanism requires to be tightly regulated in the cell to prevent a futile cycle of MglA GTP hydrolysis. But this mechanism of regulation of MglB activity in light of cell polarity reversal is not clearly understood. Interestingly, this was in contrast to another report of MglB acting only as a GAP and not as a GEF for MglA^12^. A complex of proteins comprising the response regulator, RomR along with its partner RomX was established as the GEF for MglA ^14^. Recently an activator of MglB, namely RomY was identified which further stimulates the GAP activity of MglB ^13^. MglB also acts as GAP for SofG, another small Ras-like GTPase involved in M*yxococcus xanthus* motility^15^.

The structure of the MglA-MglB complex revealed that a homodimer of MglB interacts with one molecule of MglA (MglA: MglB = 1:2) resulting in an asymmetric interaction between the two molecules^10–12^. In the asymmetric complex, the two protomers of MglB (hereafter referred to as MglB_1_ and MglB_2_) interact with different residues of the MglA monomer (Fig. 1A). Binding of MglB reorients the Switch-I and Switch-II loops of MglA. As a result of this conformational change, Thr-54, which completes the Mg^2+^ coordination in MglA, and Arg-53, which is the catalytic residue near the nucleotide-binding pocket, are optimally positioned for hydrolysis. This is a mechanism of indirect GAP activity^10,11,16^, where the GAP does not provide any of the active site residues, but functions by orienting the active site residues of the GTPase.

**Figure 1:**
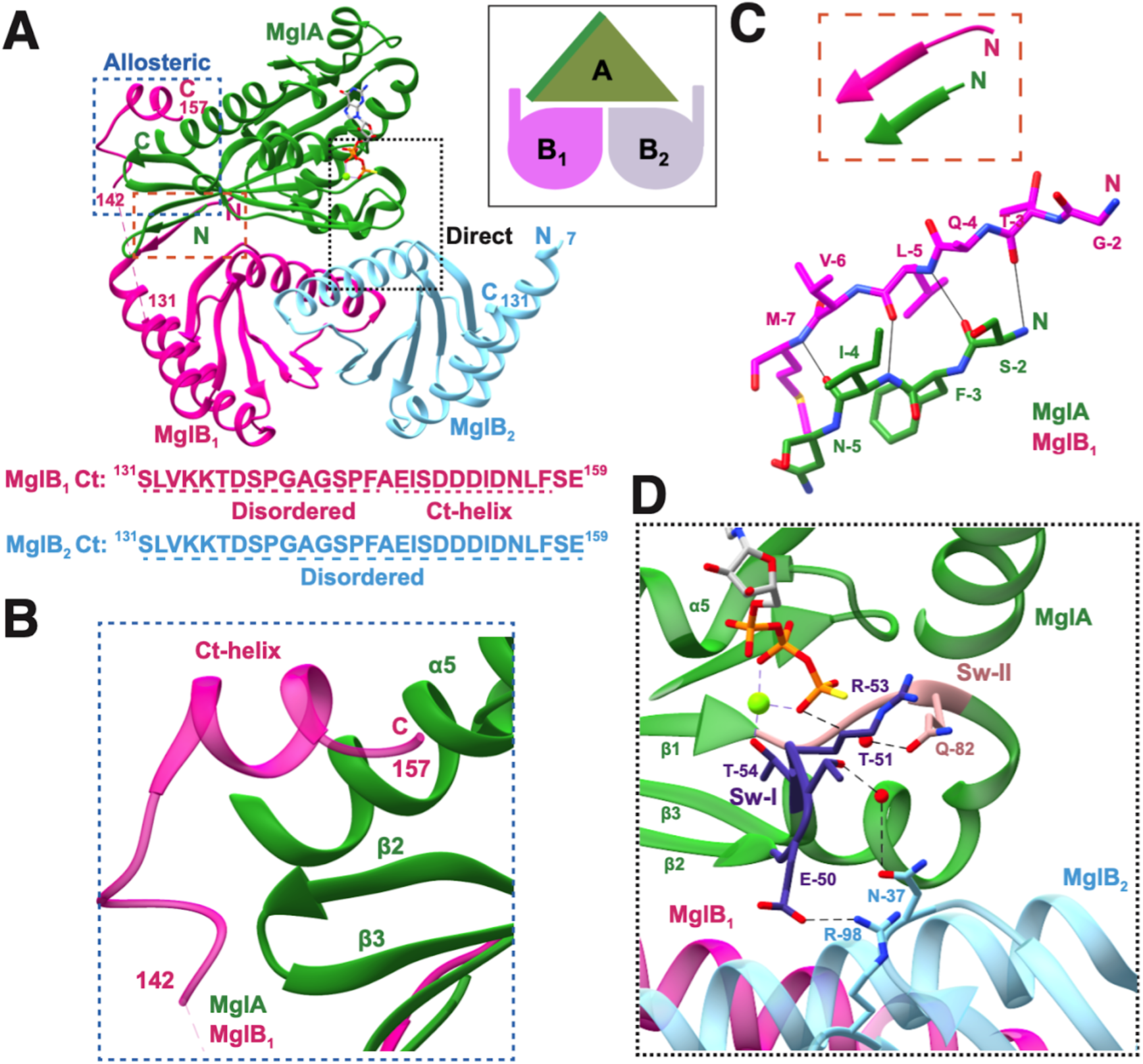
Asymmetric features of MglAB complex. **A.** Structure of the *M. xanthus* MglA-MglB complex (PDB ID: 6IZW) (MglA in green, MglB_1_ in magenta and MglB_2_ in light blue) highlighting the MglB_1_ Ct-helix interaction with MglA (blue dotted box) labelled as allosteric interaction and MglB_2_ Ct-helix interaction with MglA nucleotide-binding pocket (black dotted box) labelled as direct interaction. The residues at the C and N-termini (‘C’ and ‘N’ respectively) and the breaks of protein chains (residue numbers) are labelled for all the chains. The amino acid sequence of the Ct-region of the two MglB protomers is shown below with the disordered residues in the crystal structure and the Ct-helix sequence labelled. In the inset, a schematic representation, as depicted in later figures, of the MglAB complex is shown. **B.** Interaction of MglB_1_ Ct-helix (magenta) (start and end residue numbers labelled) with MglA (green) helix binding pocket. MglA α_5_ helix and the β_2_-β_3_ loop have been labelled. **C.** Interaction between MglB_1_ N-terminal β-strand and β_0_-strand of MglA which extends the MglA β-sheet constituting the central Ras fold. **D.** Interaction between Asn-37 and Arg-98 residues of the MglB_2_ protomer with Switch-I region (dark purple) of MglA. Switch-I and Switch-II residues are labelled in dark purple and peach respectively.

Another feature of the asymmetric MglA-MglB complex (PDB ID: 6IZW) is that the C-terminal α-helix (Ct-helix) of MglB_1_ binds in a pocket formed by the α_5_ helix and the extended β_2_-β_3_ loop of MglA (Fig. 1B). According to our earlier observations, the Ct-helix of MglB is the GEF-active region^11^. A C-terminal truncated construct of MglB (last 20 amino acids of MglB truncated, hereafter termed as MglB^ΔCt^)^11^ interacted only with MglA-GTP state, compared to the full-length MglB which has an affinity for MglA also in the GDP bound state. It was also observed that the Ct-helix of MglB is responsible for the nucleotide exchange activity on MglA^11^, by stimulating the release of GDP. In the crystal structure, the Ct-helix is ordered in the MglB_1_ monomer only. The corresponding residues in MglB_2_ are not ordered in the crystal structure, indicating that they are flexible.

The N-terminal residues of the two protomers contribute to an additional asymmetric feature. In MglB_1_, the first helix of the Rbl/LC7 fold extends from Glu-11 whereas, in MglB_2_, the helix initiates from Tyr-8. The preceding residues 1–7 are disordered in MglB_2_ (Fig. 1A). However, the residues 2-7 of MglB_1_ constitute a β-strand that forms hydrogen bonds with the β_0_ strand of MglA which extends the central β-sheet in the MglA fold (Fig. 1C). These residues of MglB thus form part of its binding interface with MglA. Hence, it is interesting to find whether this strand plays a role in orienting MglB optimally with respect to MglA.

The existence of dual GAP and GEF activities by MglB necessitates that the activities need to be tightly regulated to control the ON and OFF states of MglA in the cell. Comparison of the crystal structures of MglA-GTP equivalent states with and without MglB (PDB ID: 6IZW, 6H17) shows that Switch-I takes a different conformation in the absence of MglB ^11,12^. Asn-37 and Arg-98 of MglB_2_ interact with Switch-I residues of MglA in the MglB-bound conformation and it is plausible that the interactions might specifically regulate MglB GAP activity (Fig. 1D).

The most common mechanism of GEF activity for small Ras-like GTPases involves direct binding of GEF residues at the vicinity of the nucleotide-binding pocket, such that nucleotide and/or Mg^2+^ are evicted from the pocket^17^. The disordered C-terminal end of the MglB_2_ protomer lies close to the nucleotide-binding pocket of MglA and could drive nucleotide exchange by directly interacting with the switch regions of MglA. Interestingly, the C-terminal end (last 20 residues) of MglB contains a stretch of conserved negatively charged residues (Fig. 1A), which might compete with the binding of phosphate moieties of the nucleotide to MglA. This plausible mechanism of direct GEF activity is very similar to the TRAPPI complex, where a C-terminal extension of Bet3p, which contains conserved glutamate, stretches to the nucleotide-binding pocket of the Rab GTPase Ypt1p^18^.

Alternatively, the ordered Ct-helix from the MglB_1_ protomer in the crystal structure of the MglAB complex (6IZW), which lies opposite to the MglA nucleotide-binding pocket, suggests a unique allostery-based mechanism that can indirectly regulate nucleotide exchange by MglA. A comparison of MglA-GDP (PDB ID: 5YMX) bound and MglAB-GMPPNP (PDB ID: 6IZW) bound structures reveals a registry shift of two amino acids and a unique flipping of the β_2_ strand in MglA (β-screw movement), which exposes the hydrophobic side chains towards the MglB binding interface^10–12^. If the interaction of the Ct-helix of the MglB_1_ protomer stabilizes the conformation of MglA with the flipped β_2_-strand, this might favour MglA to bind GTP over GDP, thereby resulting in nucleotide exchange ^12^. The flipped conformation also orients Asp-58, the Walker B residue that assists Mg^2+^ coordination in the GTP-bound state^19^.

For a mechanistic understanding of GEF action, it is critical to decipher whether the C-terminal region of MglB_1_ or MglB_2_ drives nucleotide exchange by MglA. With this goal, we successfully designed and purified, and biochemically characterized complexes of MglAB with C-terminal residues deleted from either MglB_1_ or MglB_2_. Consequently, we establish that the Ct-helix from MglB_1_ allosterically drives nucleotide exchange by MglA, while the C-terminal residues from MglB_2_ are not essential for GEF activity. The observation highlights a novel strategy of GEF function based on allosteric control of a conformation favourable for GTP binding. Such a mechanism of regulation by targeting the β_2_-β_3_ loop of small Ras-like GTPases is potentially found in the GEF action of many prominent small Ras-like GTPase families such as Rag and Arf GTPases, which share many common features with the MglA family of Ras-like GTPases^19^. Furthermore, truncating the N-terminal β-strand of MglB significantly reduces the GEF activity of MglB, keeping the GAP activity intact. This shows that the asymmetric interaction of the N-terminal β−strand of MglB_1_ with the MglA central β-sheet is important for orienting the MglB Ct helix. On the other hand, mutations of Asn-37 and Arg-98 to alanines (MglB^N37R98A^) keep the GEF activity intact. Instead, it reduces the GAP activity of MglB as these residues are involved in sterically orienting the switch loops of MglA to drive GTP hydrolysis. Hence, we attribute the GAP activity of MglB to the MglB dimeric interface consisting predominantly of the MglB_2_ protomer. Therefore, our study functionally dissects the contributing factors to the contrasting GAP and GEF activities performed by MglB.

## Results

### Design of the linked constructs for a functional asymmetric MglAB complex

To delineate the functional asymmetry between the two MglB protomers, it was essential to generate an asymmetric dimer of MglB, in which only one of the MglB protomers (either MglB_1_ or MglB_2_) possessed the C-terminal residues. Because the asymmetry in MglB dimer arises only upon interaction with MglA, we designed a construct linking the C-terminus of MglA with the N-terminus of MglB (Fig. 2A). The C-terminus of MglA was proximal to the N-terminus of MglB_1_ protomer. Hence, we predicted that a short linker between MglA and MglB will suffice to form a functional MglA-MglB_1_ linked construct (Fig. 2A). The amino acid linker between MglA and MglB was chosen such that it is flexible enough to allow the functional complex formation but sufficiently rigid to disallow excessive free rotation between MglA and MglB. Hence, a Gly-Ser linker (GS) between MglA and MglB was designed based on the positions of the C-terminal and N-terminal ends of MglA and MglB_1_ respectively in the crystal structure of the MglAB complex (PDB ID: 6IZW; Fig. 2A) and the crystal packing in MglA-GDP structure, where the C-terminal hexahistidine tag formed a β−strand which continued the central β-sheet of MglA (PDB ID: 5YMX; Fig. 2A).

**Figure 2:**
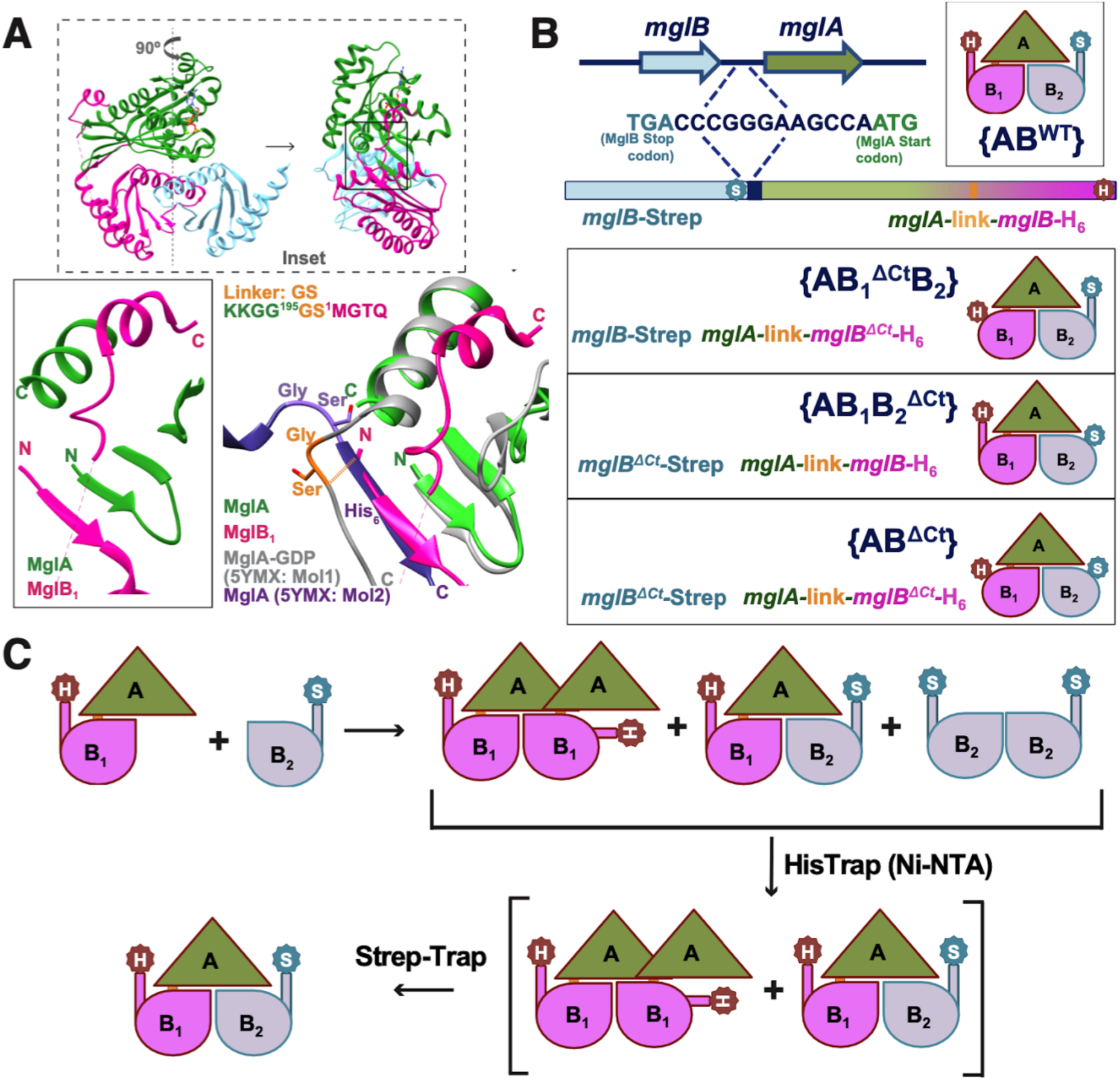
Design of the linked constructs. **A.** Orientation of N-and C-termini of MglA and MglB led to the design of a linked construct. (Insets) Structure of the *M. xanthus* MglA MglB complex (PDB ID: 6IZW; top; box in dashed outline) rotated sideways and zoomed in (bottom left; box in solid outline) to highlight the respective termini positions. A short Gly-Ser linker (light orange) between the C-terminus of MglA and the N-terminus of MglB_1_ will serve to connect MglA and MglB. The position of a similar Gly-Ser linker from the C-terminal end of MglA to its C-terminal hexahistidine tag (purple; as observed in the crystal packing of MglA-GDP structure PDB ID: 5YMX) superposed with the N-terminal β−strand of MglB_1_ is shown to demonstrate the feasibility of interaction. **B.** Strategy for co-expressing constructs of MglB_2_ (grey) and MglA-link-MglB_1_ (green-pink) (MglA: green; Linker: yellow; MglB_1_: pink) with an intervening DNA sequence from the operon (dark blue). The possible combinations of complexes thus formed are shown. The linked constructs include {AB^WT^} and those with respective truncations of MglB_1_ and MglB_2_ C-terminal residues ({AB_1_^ΔCt^B_2_}, {AB_1_B_2_^ΔCt^} and {AB^ΔCt^} respectively). MglB_2_ (outlined in blue) and MglA-link-MglB_1_ (outlined in dark red) are tagged with Strep (S: WSHPQFEK) and hexahistidine tag (H) respectively. **C.** Strategy of purification of the asymmetric complexes through sequential affinity chromatography steps (Ni-NTA and StrepTrap).

Purification from expression trials of the MglA-link-MglB_1_ construct, without co-expression with MglB, was unsuccessful because we could obtain only a negligible amount in the soluble fraction. This is possible because a dimeric MglA-link-MglB_1_ might be in negligible amounts, resulting in an exposed MglB dimer interface for the majority of the molecules. Hence, we adopted a co-expression strategy of MglA-link-MglB_1_ and MglB. Co-expression from a single plasmid was carried out where MglB_2_ and MglA-link-MglB_1_ respectively were expressed as an operon, designed based on the *mglBA* operon in *Myxococcus xanthus* genome (Fig. 2B; See Methods). In a co-expression system, an MglB monomer could interact with the exposed MglB dimer interface of the MglA-link-MglB_1_ construct. Henceforth, a complex containing a linked MglA-link-MglB_1_ construct is represented by an enclosure within ‘{’ & ‘}’ (for example, {AB^WT^}; Fig 2B), while the absence of ‘{’ and ‘}’ represents a complex formed by mixing purified batches of the different proteins (eg. AB^WT^).

MglA-link-MglB_1_ and MglB_2_ were designed with a C-terminal hexahistidine tag (GSHHHHHH) and a Strep-tag (WSHPQFEK), respectively, which could enable the purification of the linked complexes by sequential affinity chromatography (Ni-NTA and StrepTrap respectively) (Fig. 2C)^20^. This purification strategy was further applied for the C-terminal truncated constructs of MglB_1_, and MglB_2_, respectively, to generate the asymmetric complexes comprising MglB with and without the C-terminal residues ({AB_1_^ΔCt^B_2_}, {AB_1_B_2_^ΔCt^}; Fig. 2C). {AB^WT^}, a linked construct of wild type MglB with MglA, and {AB^ΔCt^}, a linked construct of C-terminal truncated MglB with MglA, functioned as controls. Any dissociation of the complexes would lead to the elution of MglB dimer as a separate fraction from the Strep-Trap column due to the higher affinity (MglB dimer contains two Strep-tags, compared to the linked complex with one Strep-tag).

### Sequential affinity chromatography yielded asymmetric linked complexes

{AB^WT^} was purified by sequential affinity-based chromatography (Ni-NTA followed by StrepTrap) which ensured successful isolation of the complex (Fig. 3A). For {AB^WT^}, the majority of the linked complex eluted from the StrepTrap column in the initial injection volumes of D-desthiobiotin (fraction I in Fig. 3A), with a negligible amount of MglB dimers (Fig. 3A) at later fractions (fraction II; MglB dimers will have higher affinity to the column due to the presence of two Strep-tags per dimer). Consequently, we proceeded with purifying all the other linked constructs with the respective C-terminal deletions (Fig. 3B-E). {AB_1_B_2_^ΔCt^} behaved similarly to {AB^WT^} through Ni-NTA and StrepTrap columns (Fig. 3B). Interestingly, fraction II of elution for {AB_1_^ΔCt^B_2_} (Fig. 3C) and {AB^ΔCt^} purifications (Fig. 3D) contained more prominent bands corresponding to MglB_2_ and bands of MglA-link-MglB_1_, suggesting a possibility of shuffling of protomers following Ni-NTA elution.

**Figure 3:**
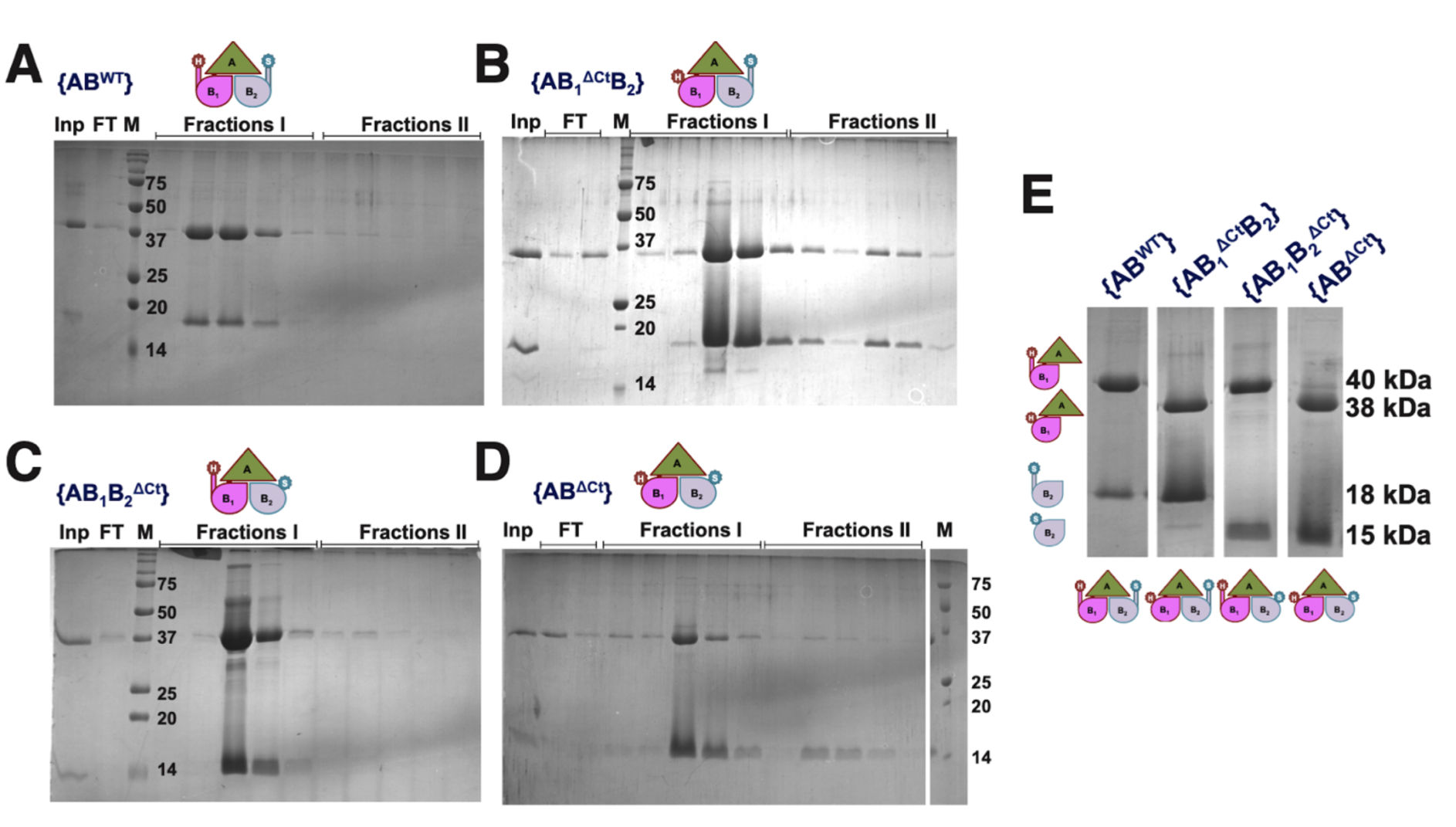
Purification of linked complexes. **A, B, C, D.** SDS-PAGE showing the StrepTrap elution profiles of the linked complexes of {AB^WT^}, {AB_1_^ΔCt^B_2_}, {AB_1_B_2_^ΔCt^} and {AB^ΔCt^} respectively. The first lane labelled as Inp is the load (obtained after Ni-NTA purification), and the lane labelled FT represents flow through (unbound protein). M is the protein ladder labelled with the molecular weights (in kDa). Fractions I and II represent the 5 eluted fractions of 1 ml each from each injection of 5 ml of 2.5 mM desthiobiotin containing elution buffer. The major constructs eluted as ‘fractions I’ are represented via the schematic. **E.** SDS PAGE showing the purified complexes from the linked constructs (cropped images of single lanes from figure panels A – D compiled for ease of comparison). The upper band corresponds to MglA-link-MglB_1_ and the lower band is of MglB_2_. Corresponding schematics of each construct are shown on the left and top of the lanes, and the sizes are labelled on the right.

The presence of C-terminal residues of MglB contributes to an aberrant movement in SEC^11^, and hence the molecular weights of the elution peaks in SEC were further confirmed using size exclusion chromatography coupled with multi-angle light scattering studies (SEC-MALS; Fig. 4A,4B). The single peak of {AB^WT^} (peak 1 eluted at 14.2 ml; Fig. 4A) corresponded to a molar mass of around 53 kDa, close to the expected mass of the complex (tabulated in Fig. 4C). The homogenous single peak of {AB_1_B_2_^ΔCt^} (peak 1 eluted at 14.9 ml) corresponded to an average molar mass of 56 kDa (Fig 4A,4C). The difference in elution volume with respect to {AB^WT^} despite the small difference in molecular weight could be attributed to the absence of aberrant movement without the MglB_2_ Ct-helix in {AB_1_B_2_^ΔCt^}.

**Figure 4:**
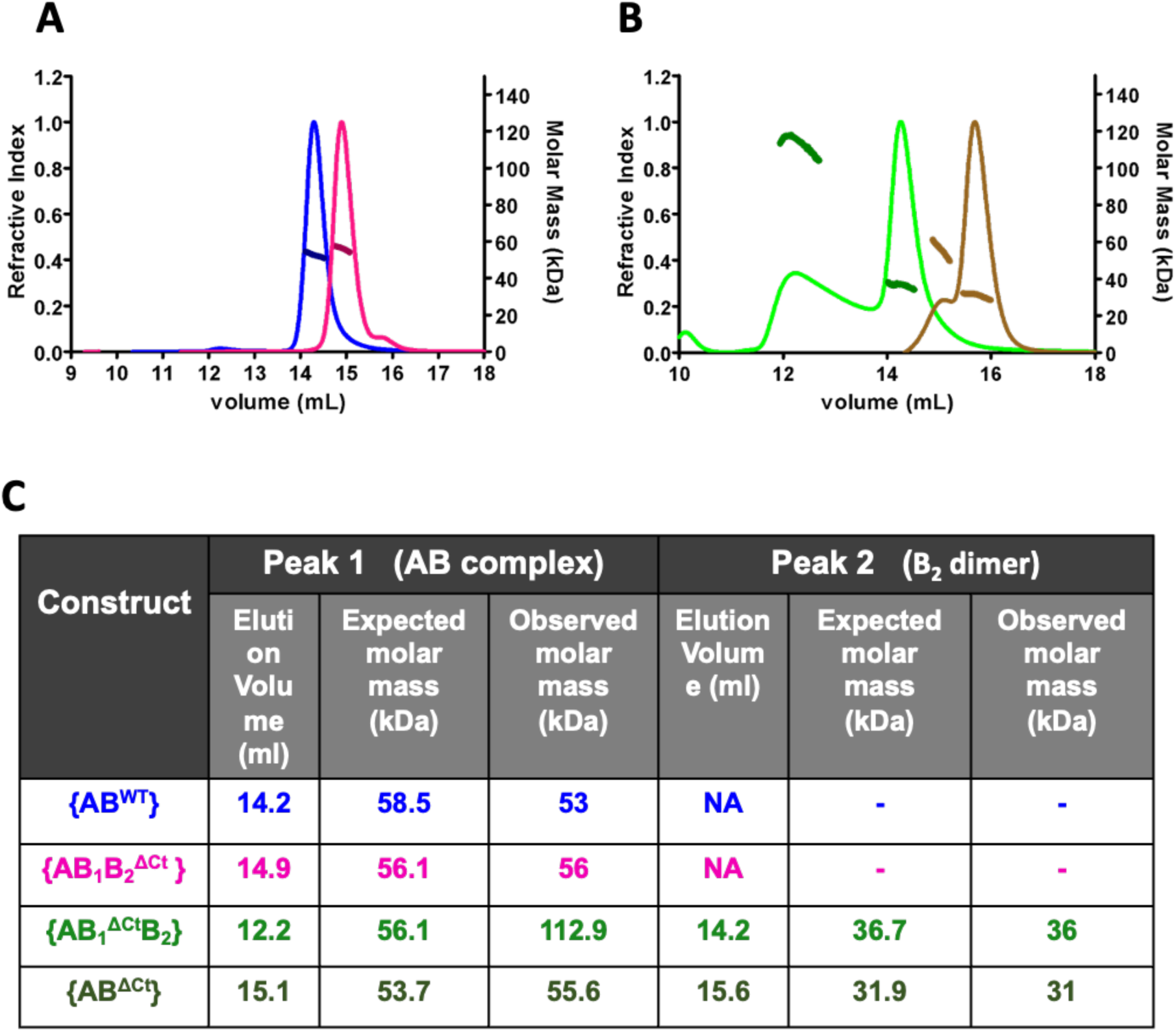
The C-terminal residues of MglB contribute to the differential stability of the linked complexes. **A, B** SEC-MALS (Superdex-200 profiles for SEC carried out at room temperature ∼ 25 °C) for the linked constructs: {AB^WT^} blue and {AB_1_B_2_^ΔCt^} magenta in panel A and {AB_1_^ΔCt^B_2_} green and {AB^ΔCt^} brown in panel B. The left and the right y axes represent the refractive index and molar mass (kDa) respectively for both plots. Molar masses are represented by thicker lines as compared to refractive indices. **C.** Table enlisting the elution volumes and corresponding average molar masses (expected and observed; Peaks 1 and 2 correspond to AB_1_B_2_ complex and B_2_ dimer, respectively as labelled in panels A and B) for the linked complexes as concluded from SEC-MALS.

However, the purified {AB_1_^ΔCt^B_2_} complex separated into two peaks in SEC (Fig. 4B). For {AB_1_^ΔCt^B_2_}, the first peak (peak 1; eluted at 12.2 ml; Fig. 4B) corresponded to a molar mass of 112 kDa which interestingly corresponded to the mass of a dimeric {AB_1_^ΔCt^B_2_} complex. However, the peak was not symmetric and showed a long tail till the second peak (peak 2; eluted at 14.2 ml; Fig. 4B) correlating with a full-length MglB dimer (36 kDa) (Fig. 4C). This tailing nature of the peak corresponded to a decreasing trend in the molar mass distribution, which could not be reliably estimated beyond 12.6 ml. This probably indicated the presence of a heterogeneous mix of monomeric and dimeric {AB_1_^ΔCt^B_2_} complexes. The dissociation of protomers was higher in the case of {AB^ΔCt^} which had a higher intensity of the MglB^ΔCt^ dimer peak (labelled as peak 2; eluted at 15.6 ml) and a smaller peak corresponding to {AB^ΔCt^} (peak 1; eluted at 15.1 ml with a molar mass of approximately 55 kDa) (Fig. 4B,4C). Thus, the sequential affinity purification and SEC results of the linked complexes showed that the presence of the MglB_1_ helix is critical for a stable complex consisting of MglA-link-MglB_1_ and MglB_2_. This is consistent with our previous observation of interaction between GDP-bound MglA and MglB only in the presence of the MglB C-terminal residues^11^. From the observation of a stable complex only when MglB_1_ C-terminal helix is present, this stabilization effect is probably a contribution from the MglB_1_ C-terminal helix while the absence of it destabilises the complex.

The presence of elution peaks corresponding to the molecular weight of MglB_2_ dimers (either MglB or MglB^ΔCt^ in {AB_1_^ΔCt^B_2_} and {AB^ΔCt^} purification, respectively) proved that in the absence of the MglB_1_ helix, dissociation of the linked MglAB complex resulted in free MglB_2_ protomers from the complex. The incompatibility and steric hindrance of MglB dimer, especially MglB_2_ protomer, with the MglA-GDP conformation, might result in destabilization of the dimer interface and its dissociation, which consequently might form MglB dimers or trans oligomers (as observed in {AB_1_^ΔCt^B_2_} complex) with MglA-link-MglB_1_ in solution. Hence, we conclude that the MglB_1_ Ct-helix interaction with MglA is essential for the formation of the stable and functionally relevant linked MglAB complexes.

### The absence of MglB_2_ Ct-helix does not affect the GTPase activity of MglA

Following the purification of the linked MglAB complexes, the GTPase activity of the linked complexes was quantified using an NADH-coupled enzymatic assay to monitor GDP release. It was observed that {AB^WT^} showed comparable *k_obs_* values to the unlinked AB^WT^ complex which indicated the formation of a functional linked complex (Fig. 5A,Table 1). Interestingly, {AB_1_B_2_^ΔCt^} exhibited GTP hydrolysis rates comparable to that of the full-length complexes (linked and unlinked). This shows the C-terminal residues of the MglB_2_ protomer did not play a major role in increasing the net GTPase activity of MglA (Fig. 5A,Table 1). The least activity was observed for {AB_1_^ΔCt^B_2_} and {AB^ΔCt^} complexes both of which had the MglB_1_ Ct-helix truncated. Incidentally, the activity was equivalent to that displayed by the complex formed by MglA and MglB^ΔCt^ (AB^ΔCt^) where the Ct-helix was absent in the MglB dimer. Comparable activities of unlinked MglA and MglB^ΔCt^ complex and linked {AB_1_^ΔCt^B_2_} and {AB^ΔCt^} suggest that the concentrations of the MglA domain is similar in all these cases and the reduced activity is probably due to the absence of the MglB_1_ Ct-helix. Hence the reduced activity for {AB_1_^ΔCt^B_2_} indicates that the MglB_2_ C-terminal residues did not contribute to an increase in GTPase activity of MglA. Thus, the deletion of MglB_2_ C-terminal residues did not significantly affect MglA GTP hydrolysis, while the presence of the MglB_1_ helix was preferentially critical for accelerating the rate.

**Figure 5:**
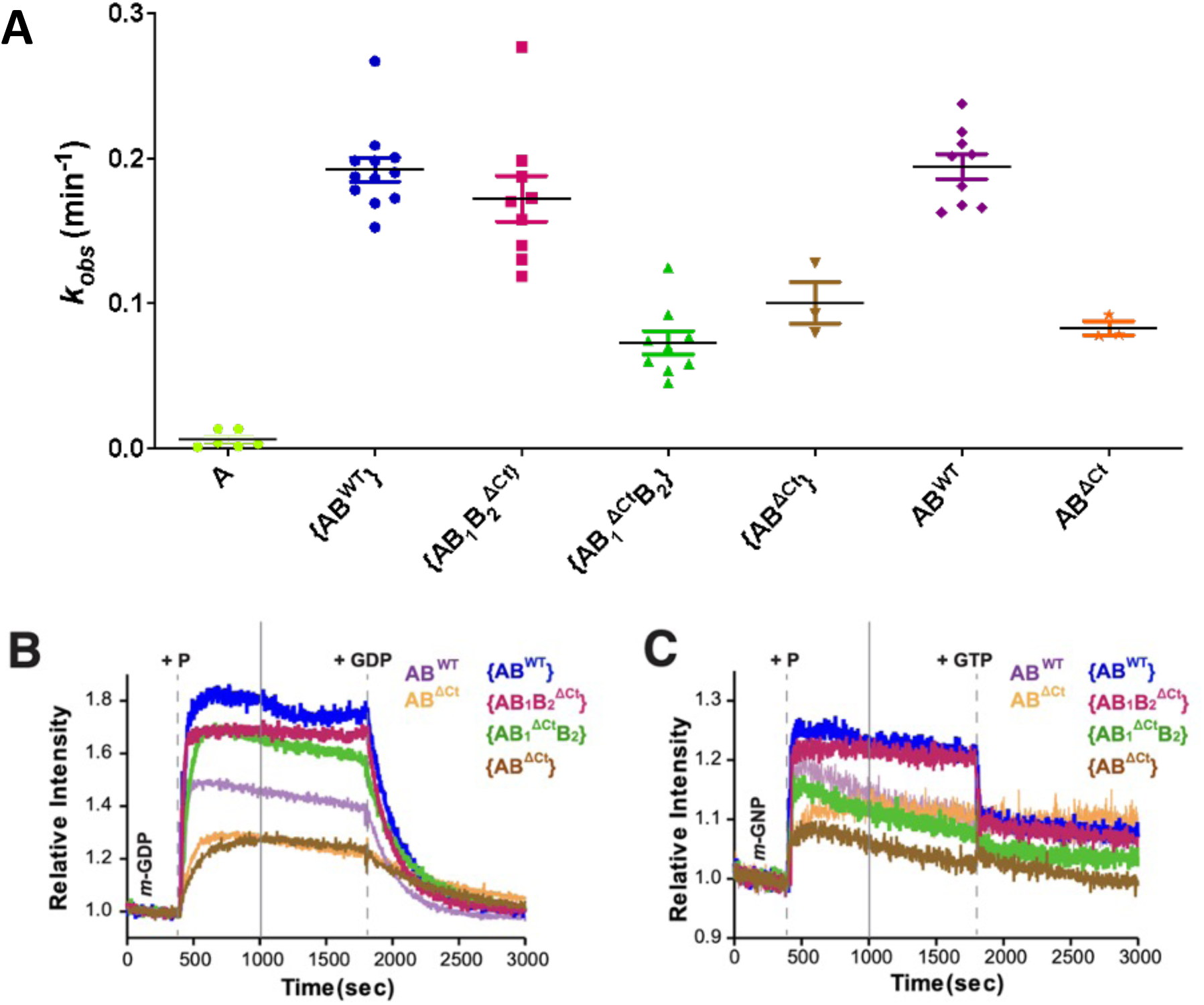
Constructs with MglB_1_ helix stimulate higher GTP hydrolysis rates of MglA and nucleotide exchange, **A.** Comparison of *k_obs_* values of {AB^WT^} (blue), {AB_1_^ΔCt^B_2_} (green), {AB_1_B_2_^ΔCt^} (magenta), {AB^ΔCt^} (brown), unlinked AB^WT^ (purple), AB^ΔCt^ (coral). The mean and 95% confidence intervals are shown by long and short horizontal lines, respectively for each sample. **B.** Kinetics of increase in *mant-*GDP fluorescence (region labelled as *m-*GDP) upon adding {AB^WT^} (blue), {AB_1_^ΔCt^B_2_} (green), {AB_1_B_2_^ΔCt^} (magenta), {AB^ΔCt^} (brown), AB^WT^ (purple), AB^ΔCt^ (coral), at 400 seconds (marked by the dashed line labelled ‘+P’; Stage I), followed by a competition of *mant-*GDP by adding excess unlabelled GDP at 1800 seconds (marked by the dashed line labelled ‘+GDP’; Stage II). **C.** Kinetics of increase in *m-*GMPPNP fluorescence (region labelled as *m-*GNP) upon adding {AB^WT^} (blue), {AB_1_^ΔCt^B_2_} (green), {AB_1_B_2_^ΔCt^} (magenta), {AB^ΔCt^} (brown), AB^WT^ (purple), AB^ΔCt^ (coral), at 400 seconds (marked by the dashed line labelled ‘+P’; Stage I), followed by a competition of *mant-*GMPPNP by adding excess unlabelled GTP at 1800 seconds (marked by the dashed line labelled ‘+GTP’; Stage II). The phases I and II represent association and dissociation for kon and koff estimation respectively (tabulated in Table 2A), and are demarcated by solid lines.

**Table 1:** List of observed *k_cat_* (*k_obs_* in min^-1^, *k_obs_* here is the amount of GDP released per minute per µM enzyme, for details, see Materials & Methods)

| Construct | $k_{obs}$ ($\text{min}^{-1}$) |
| --- | --- |
| MglA | $0.009 \pm 0.003$ (n=3) |
| MglB | $0.02 \pm 0.002$ (n=3) |
| {AB<sup>Wt</sup>} | $0.19 \pm 0.008$ (n=13) |
| {AB<sub>1</sub><sup>$\Delta\text{Ct}$</sup>B<sub>2</sub>} | $0.07 \pm 0.008$ (n=9) |
| {AB<sub>1</sub>B<sub>2</sub><sup>$\Delta\text{Ct}$</sup>} | $0.16 \pm 0.016$ (n=10) |
| {AB<sup>$\Delta\text{Ct}$</sup>} | $0.10 \pm 0.014$ (n=3) |
| AB<sup>Wt</sup> | $0.19 \pm 0.02$ (n=7) |
| AB<sup>$\Delta\text{Ct}$</sup> | $0.08 \pm 0.005$ (n=3) |
| MglAB<sup>$\Delta\text{N}$</sup> | $0.08 \pm 0.006$ (n=4) |
| MglAB<sup>N37R98A</sup> | $0.11 \pm 0.004$ (n=4) |

**Table 2:**
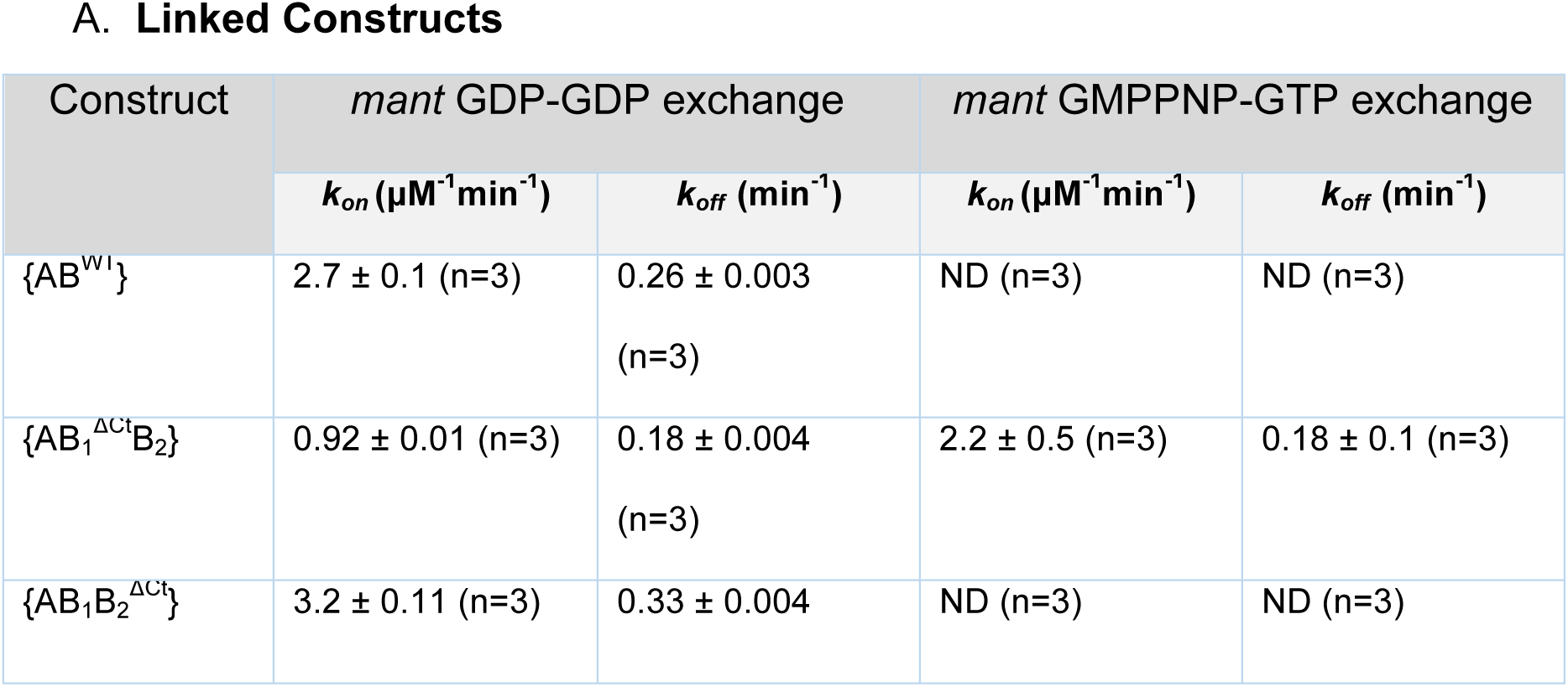
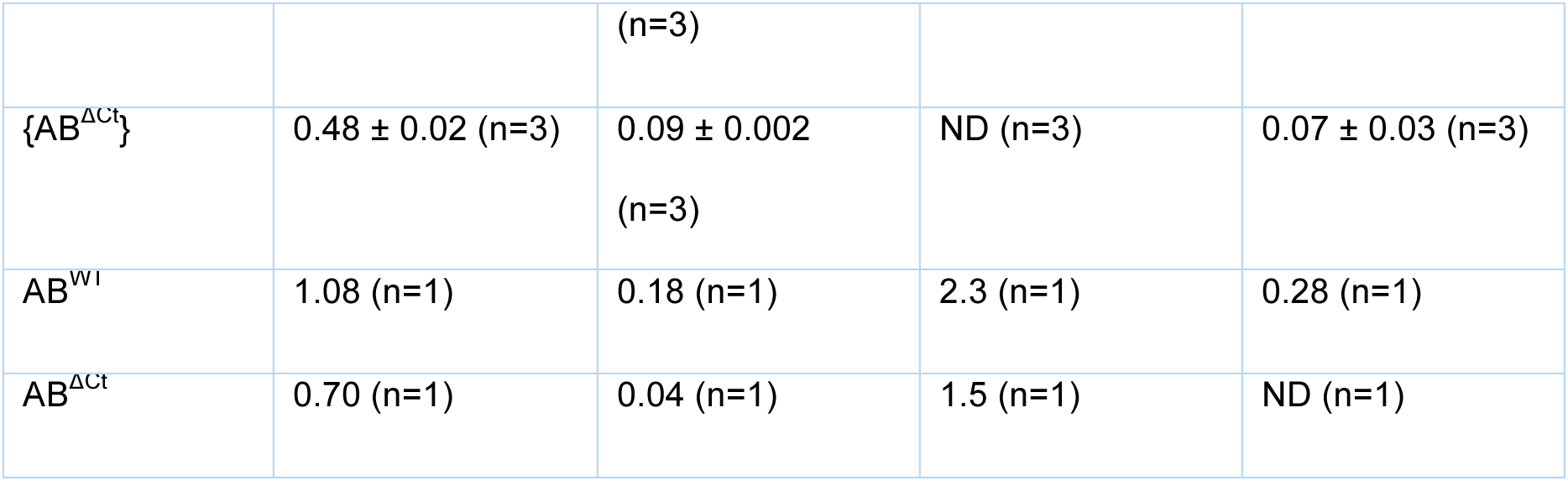
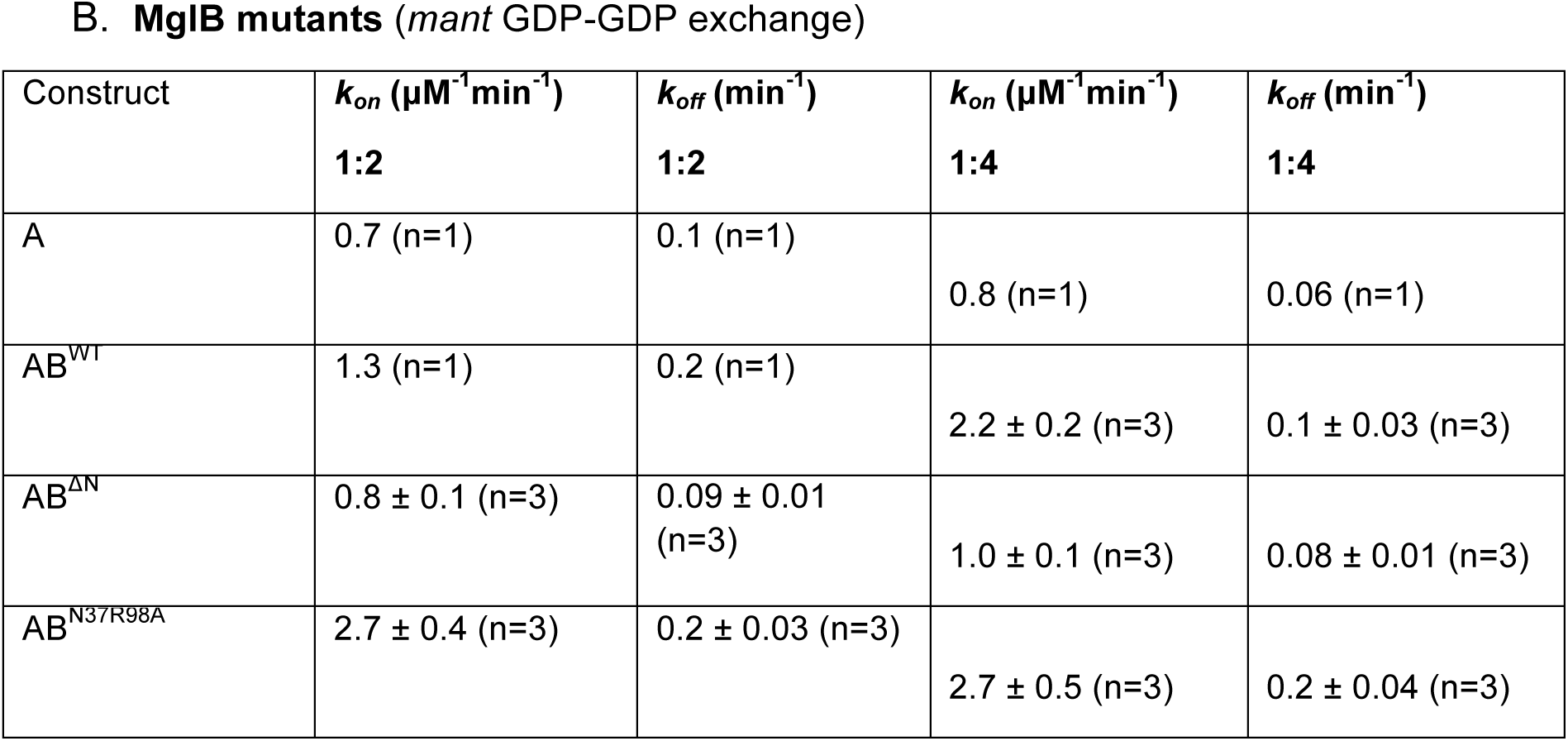
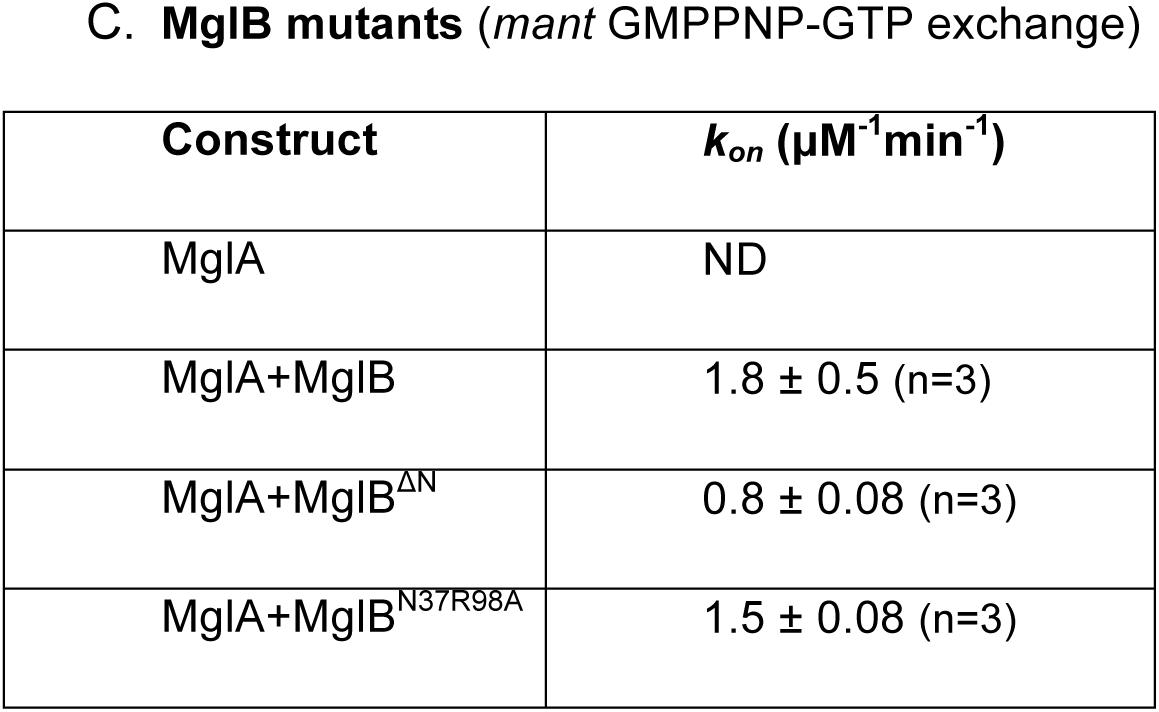
List of rates of association and dissociation (*k_on_*(µM^-1^min^-1^) and *k_off_* (min^-1^) respectively).

### MglB_1_ Ct-helix is sufficient for nucleotide exchange in MglA

Next, we performed nucleotide exchange assays with fluorescently labelled *mant*-GDP/GMPPNP. We first investigated *mant*-GDP binding to the linked complexes (labelled as Stage I) followed by its dissociation (labelled as Stage II) with excess unlabelled GDP (Fig. 5B). {AB^WT^} and {AB_1_B_2_^ΔCt^} showed very fast association with *mant*-GDP (*k_on_* = 2.7 µM^-1^min^-1^ and 3.2 µM^-1^min^-1^, respectively). The association rates of {AB_1_^ΔCt^B_2_} were slower, similar to the unlinked AB^WT^ complex (*k_on_* _=_ 0.9 µM^−1^min^-1^). {AB^WT^} and {AB_1_B_2_^ΔCt^} showed higher rates of exchange than unlinked AB^WT^ (Table 2A), probably owing to the tethering of MglB to MglA which increases the probability of the MglB Ct-helix association with MglA. {AB_1_^ΔCt^B_2_} shows exchange with a slower *k_on_* (0.92 µM^-1^min^-1^) than AB^WT^. This exchange activity is probably due to the combined effect of the presence of a linked MglB_1_ and from the *trans* activity by the full-length MglB_2_ dimer, formed post-dissociation of the complex as indicated by the SEC-MALS study. The slowest *k_on_* of 0.45 µM^-1^min^-1^ was exhibited by {AB^ΔCt^} because both the linked MglB_1_ protomer and the MglB_2_ dimer, formed from the dissociated protomers, lacked the C-terminal residues. This was similar to what was observed in unlinked AB^ΔCt^ (0.6 µM^-1^min^-1^). The rates of exchange for dissociation of *mant*-GDP by adding unlabelled GDP (*k_off_*) were again highest for {AB^WT^} and {AB_1_B_2_^ΔCt^} (0.26 min^-1^ and 0.33 min^-1^, respectively) followed by {AB_1_^ΔCt^B_2_} (0.18 min^-1^). {AB^ΔCt^} (*k_off_* = 0.09 min^-1^) exhibited slower exchange as the exchange of GDP was not facilitated in the absence of the MglB Ct-helix, similar to unlinked AB^ΔCt^.

Similar nucleotide exchange experiments were performed with *mant*-GMPPNP followed by dissociation with excess unlabelled GTP (Fig. 5C) to further resolve the effect of the MglB Ct-helix deletions. {AB^WT^} and {AB_1_B_2_^ΔCt^} showed very fast association and dissociation rates of *mant*-GMPPNP and hence values of *k_on_* and *k_off_* could not be estimated. This could be because the linked MglB_1_ Ct-helix in both these complexes drove nucleotide exchange efficiently. {AB_1_^ΔCt^B_2_}, without a Ct-helix in the MglB_1_ protomer, exhibited slower exchange kinetics as unlinked AB^WT^ with measurable *k_on_* and *k_off_* values of 2.2 µM^-1^min^-1^ and 0.18 min^-1^, respectively. {AB^ΔCt^} exhibited exchange rates similar to unlinked AB^ΔCt^, however, due to the lack of stability of the complex we observed inconsistent signals for *mant*-GMPPNP binding across replicates. Unlike in GTPase assays (where a GTP concentration of 1 mM was used), a lower concentration of *mant*-GMPPNP (800 nM) failed to stimulate GTP binding and stabilization for this complex. Hence, we failed to obtain reliable estimates of *k_on_*. *k_off_* was negligibly low (0.07 min^-1^). All these results indicate that in {AB^WT^} and {AB_1_B_2_^ΔCt^}, the exchange of GDP with GTP is driven by the MglB_1_ Ct-helix, which interacts away from the nucleotide-binding pocket of MglA.

### N-terminal β-strand of MglB_1_ helps orient the Ct helix driving nucleotide exchange

Next, we attempted to dissect the role of the asymmetric interaction of the N-terminal region of MglB_1_ with MglA. Residues 2-7 were deleted in MglB, which constituted the β-strand in MglB_1_ that interacts with β_0_-strand of MglA. This strand might play a role in orienting MglB and facilitating optimal interaction with MglA. This construct was named MglB^ΔN^. Size exclusion chromatography confirmed that MglB^ΔN^ was well folded and formed a dimer like MglB (Fig. 6A). Next, we compared the GTPase activity of MglA in presence of MglB^ΔN^. MglA GTP hydrolysis rate in the presence of the mutant MglB was reduced as compared to that with MglB^WT^ (Fig. 6B,Table 1). *k_obs_* was similar to that with MglB^ΔCt^ (Table 1) which had only GAP and no GEF activity due to the absence of the Ct helix. We further checked the GDP exchange rates of MglA in presence of MglB^ΔN^ using fluorescently labelled *mant* GDP (Fig. 6C). We observed that the *k_on_* and *k_off_* rates (0.8 µM^-1^min^-1^ and 0.09 min^-1^, respectively) are similar to MglA alone which proved that the GEF activity of MglB^ΔN^ was abolished (Table 2B). We performed the same assay with excess concentrations of MglB mutants (MglA:MglB of 1:4) and observed similar results (Supplementary Fig 2). This validates that the reduction of the GEF activity of mutant MglB is not due to reduced affinity to MglA. We also observed GDP exchange rates using fluorescently labelled *mant* GMPPNP. MglB^ΔN^ showed a slower *k_on_* rate similar to MglB^ΔCt^ (0.8 µM^-1^min^-1^) (Table 2C). All these results showed that the deletion of the N-terminal β−strand in MglB hindered the optimal interaction of the MglB_1_ Ct helix with MglA thereby abrogating the GEF activity.

**Figure 6:**
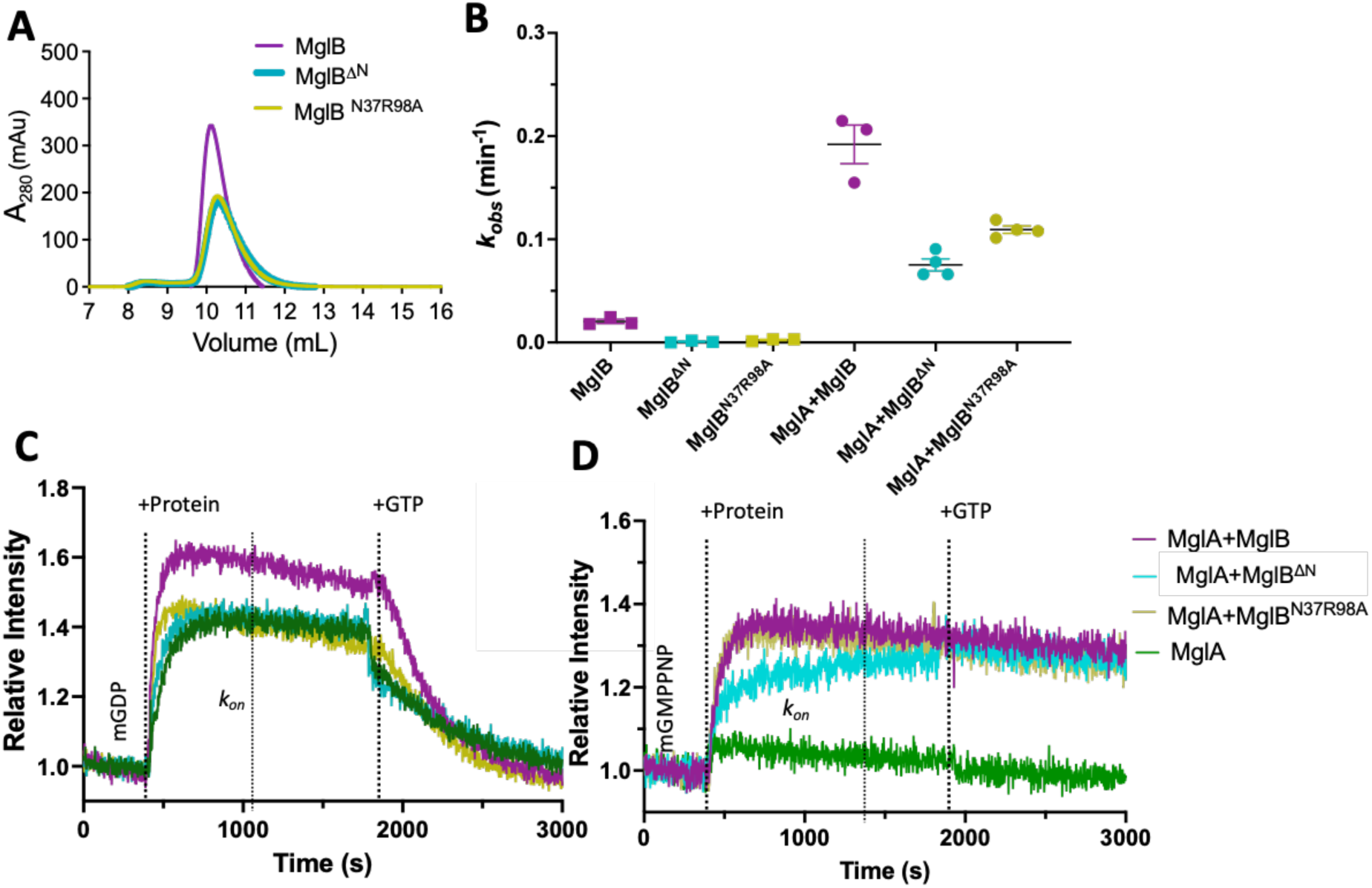
MglB_1_ N terminal β-strand and MglB_2_ Asn-37/Arg-98 are essential for MglB GEF and GAP activity, respectively. **A.** Size exclusion chromatography (Superdex-75) profiles of mutants of MglB (purple), MglB^ΔN^ (cyan) and MglB^N37A,R98A^ (ochre). **B.** Comparison of *k_obs_* values of MglB control (square) with MglB^WT^ (purple) with MglB asymmetry mutants MglB^ΔN^ (cyan) and MglB^N37R98A^ (ochre) and respective constructs with MglA (circles). The mean and 95% confidence intervals are shown by long and short horizontal lines, respectively for each sample. **C.** Kinetics of increase in *mant-*GDP fluorescence (region labelled as *m-*GDP) upon adding MglA (dark green), unlinked AB^WT^ (purple), MglA with MglB mutants, MglB^ΔN^ (cyan) and MglB^N37R98A^ (ochre) at 400 seconds (marked by the dashed line labelled ‘+P’; Stage I), followed by a competition of *mant-*GDP by adding excess unlabelled GDP at 1800 seconds (marked by the dashed line labelled ‘+GDP’; Stage II). **D.** Kinetics of increase in *mant-*GMPPNP fluorescence (region labelled as *m-* GMPPNP) upon adding MglA (dark green), unlinked AB^WT^ (purple), MglA with MglB mutants, MglB^ΔN^ (cyan) and MglB^N37R98A^ (ochre) at 400 seconds (marked by the dashed line labelled ‘+P’; Stage I), followed by a competition of *mant-*GMPPNP by adding excess unlabelled GTP at 1800 seconds (marked by the dashed line labelled ‘+GTP’; Stage II). The phases I and II represent association and dissociation for kon and koff estimation respectively (tabulated in Table 2B and 2C), and are demarcated by solid lines.

### Asn-37 and Arg-98 of MglB are essential for MglB GAP activity

It is evident from the structure of the MglA/MglB complex (GTP bound; PDB: 6IZW; Fig. 1A) that the interaction of MglB reorients Switch-I region of MglA for active hydrolysis thereby functioning as a GAP. Since the residues Asn-37 and Arg-98 of MglB_2_ interact with the Switch-I residues of MglA (Fig. 1D), these residues might be critical for Switch-I reorganization and holding it in its active conformation thereby contributing to the MglB GAP function. A double mutant was, hence, designed where we mutated Asn-37 and Arg-98 to alanines (MglB^N37R98A^), to investigate the role of these residues in activating MglA GTP hydrolysis. The mutation did not affect the folding or oligomeric status of the purified protein as inferred from size exclusion chromatography (Fig. 6A). However, the GTPase stimulation rate of MglA by MglB^N37R98A^ was slightly reduced as compared to wild-type MglB as evident from the observed GTP hydrolysis rates (Fig. 6B,Table 1). To further dissect the role of the mutations, we performed a nucleotide exchange assay with *mant* GDP and competed it out with unlabelled GDP (Fig. 6C). We observed that the *k_on_* rate of MglA with MglB^N37R98A^ (2.7 µM^-1^min^-1^) was slightly higher than with MglB (1.3 µM^-1^min^-1^) (Table 2B). However, the *k_off_* rate was similar to MglA+MglB (0.2 min^-1^). However, the *k_on_ and k_off_* rates were comparable to MglA+MglB when we performed the same assay with an excess of MglB mutants (MglA: MglB 1:4) (Supplementary Fig. 2). This showed that the GEF activity of MglB^N37R98A^ was intact. In similar assays done with *mant* GMPPNP and competition with unlabelled GTP, we observed a similar trend where the *k_on_* rate of MglA with MglB^N37R98A^ (1.5 µM^-1^min^-1^) was again similar to that observed for MglA+MglB (1.8 µM^-1^min^-1^) (Table 2C). Hence it was concluded that the interactions of Asn-37 and Arg-98 of the MglB_2_ protomer with the MglA Switch-I region were important for accelerating the MglB GAP activity. All these results highlight that the MglB dimeric interface predominantly consisting of the MglB_2_ protomer is the GAP active region of MglB whereas the Ct-helix of the MglB_1_ protomer is the GEF active region.

## Discussion

We successfully designed linked constructs of the MglAB complex with MglB Ct-helix truncations to dissect the functional asymmetry of the interaction between MglA and MglB. Our study employs a unique approach to dissect the functional basis of an asymmetric interaction of a multimeric regulator of a GTPase. We have succeeded in resolving the respective GAP and GEF active regions of the MglB, *in vitro*, which can eventually throw light into how this dual function of MglB is spatially resolved in a cell exhibiting polarity reversals.

Earlier studies showed that the Ct-helix truncated construct of MglB acts as a GAP ^10^ and the C-terminal region contributes to the GEF activity^11^. {AB_1_^ΔCt^B_2_}, which lacked the MglB_1_ Ct-helix, possessed hydrolysis rates similar to {AB^ΔCt^} where the helix was truncated in both the MglB protomers. In nucleotide exchange assays, both {AB^WT^} and {AB_1_B_2_^ΔCt^} exhibited efficient GDP and GTP exchange as both the complexes had MglB_1_ helix interacting with MglA. The accelerated rates of exchange in {AB^WT^} and {AB_1_B_2_^ΔCt^} could be attributed to the linked complex which increases the probability of the interaction of the MglB_1_ Ct-helix with the helix binding pocket of MglA. In {AB_1_^ΔCt^B_2_}, the dissociated MglB_2_ could dimerise and the Ct-helix was probably capable of interacting with MglA in *trans*. This resulted in GDP and GTP exchange rates of {AB_1_^ΔCt^B_2_} similar to unlinked MglA/MglB complex where the interaction of MglA with MglB depended on the diffusion of the two components. Hence, the comparison of {AB^WT^} and {AB_1_B_2_^ΔCt^} shows that the GEF-active component is the MglB_1_ helix that acts via the helix binding pocket on MglA which lies opposite to the nucleotide-binding pocket.

Another component of the asymmetric interaction is the β-strand of the MglB_1_, an extension of the Rbl/LC7 fold of the MgB_1_ protomer interacting with the β_0_ strand of the central β-sheet of MglA. This interaction could potentially help in maintaining a correct orientation of MglB with respect to MglA. The β_0_ strand is unique for the MglA family. Deletion of interacting β-strand in MglB does not affect the stability of MglB. However, it does reduce the GTPase rate of MglA as compared to that of MglB^WT^. The GAP activity was intact but the GEF activity of MglB^ΔN^ was affected. We compared the conformations of residues 1 to 7 in the structure of MglB only (PDB ID: 6HJM Chains A, B) and MglAB complex (PDB ID: 6IZW). MglB_1_ forms the N-terminal β-strand only in the presence of MglA. In MglB_2_, the first α-helix of Rb/LC7 is longer compared to MglB_1_. If MglB_1_ also had a continuous longer α-helix, it might hinder the interaction of the Ct-helix of MglB_1_ onto MglA (Fig. 1B). We speculate that the formation of the N-terminal β−strand of MglB_1_ and further its interaction with MglA helps to avoid clashing of N-terminal region of MglB and Ct-helix of MglB. Also, deletion of this strand probably hinders the optimal interaction of MglB with MglA thereby affecting the GEF activity of MglB.

Asn-37 and Arg-98 residue mutants of MglB were designed to dissect the role of these residues in orienting the Switch region of MglA. The double mutation reduced the GTPase stimulation of MglB. However, the nucleotide exchange activity was intact despite the mutation. Hence, these residues from MglB_2_ protomer interacts and orients the Switch-I residues of MglA promoting MglB GAP activity. MglB contains a roadblock domain (RD), which is the ancestor of the Longin domain, a characteristic domain present in GEFs of eukaryotic small Ras-like GTPases. MglB RD forms a dimer symmetrically to form an extended β-meander and forms a platform, which interacts asymmetrically with one MglA molecule^21^.

The mutation of Asn-37 and Arg-98 residues of MglB was designed to dissect the role of these residues in orienting the Switch region of MglA. The double mutation reduced the GTPase stimulation of MglB. However, the nucleotide exchange activity was intact despite the mutation. Hence, these residues from MglB_2_ protomer interacts and orients the Switch-I residues of MglA promoting MglB GAP activity. MglB contains a roadblock domain (RD), which is the ancestor of the Longin domain, a characteristic domain present in GEFs of eukaryotic small Ras-like GTPases. MglB RD forms a dimer symmetrically to form an extended β-meander and forms a platform, which interacts asymmetrically with one MglA molecule^21^. It is interesting to compare such an asymmetric interaction in the MglAB complex to that of a symmetric interaction in Rag GTPase dimer. Uniquely, Rag GTPases have an RD in *cis* towards its C-terminus, analogous to MglA-link-MglB_1_ in our study. This RD mediates the dimerization of two subunits^22^. However, the interface analogous to the interface between MglA and MglB dimer does not exist in Rag GTPases. The RD dimer forms a platform to support the respective GTPase domains such that each of them can be independently activated to a conformation facilitating GTP hydrolysis (GAP activity)^23,24^. Extensions to the RD domain functions as a GEF in eukaryotes through direct interaction at the nucleotide-binding pocket^18^, however, we show that the extension to the MglB Rbl domain functions through an independent allosteric mechanism involving the C-terminal helix (Fig. 7).

**Figure 7:**
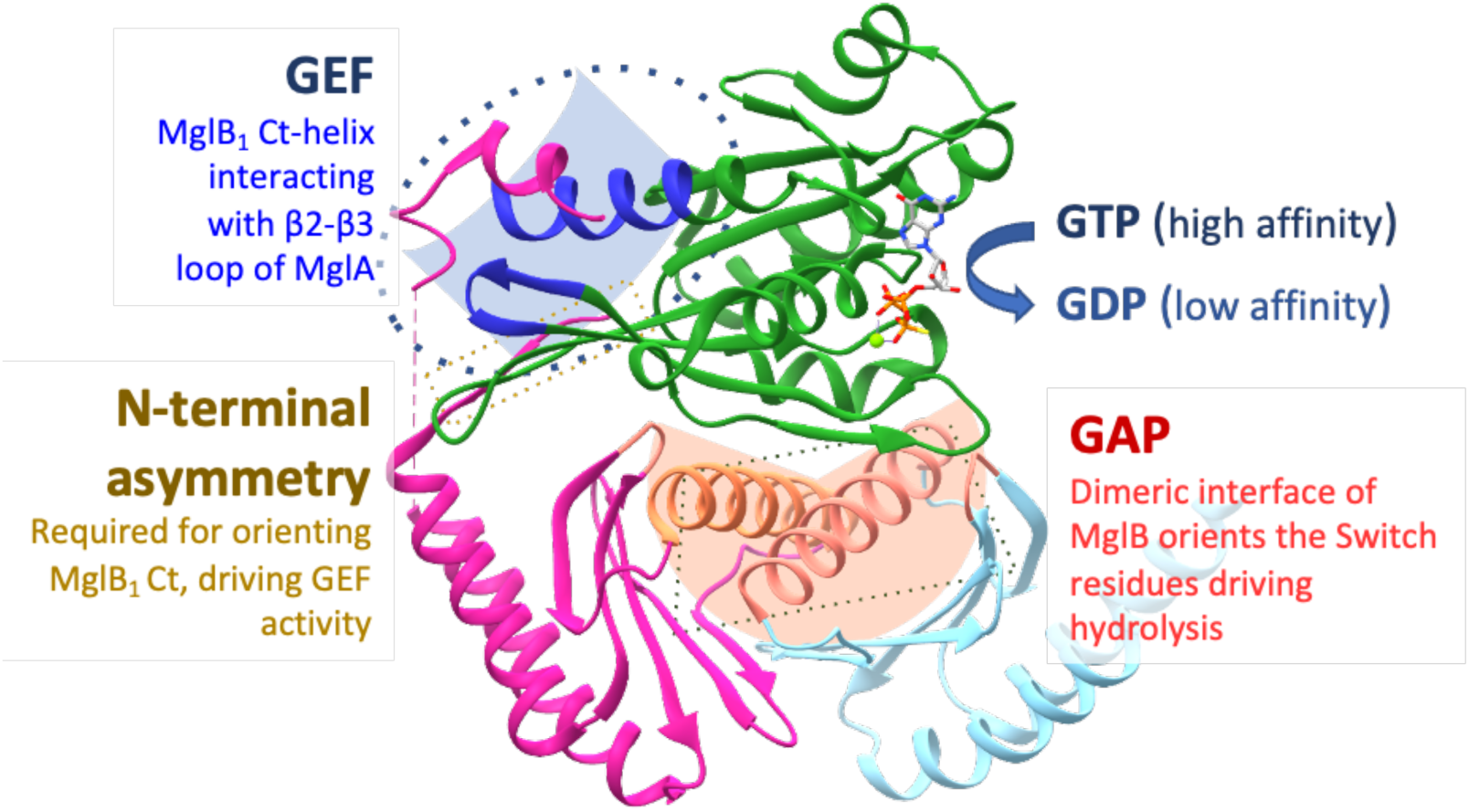
Mechanism of dual GAP and GEF activity of MglB. Asymmetric complex of MglA (green) with MglB_1_ (magenta) and MglB_2_ (light blue) with interfaces annotated with the GAP and GEF functions of MglB. C-terminal helix of MglB_1_ interacts with the MglA helix binding pocket (blue) imparting the GEF activity. The absence of the N-terminal region of MglB_1_ (yellow) precludes MglB_1_ Ct helix interaction. The dimeric interface of MglB (orange) stabilizes MglA and promotes the GAP function.

The recently identified module comprising RomR-RomX has been shown to act as a GEF for MglA^14^. It recruits MglA-GTP to the leading pole of the cell thereby establishing the front-rear polarity. Interestingly, both MglB and RomR-RomX localise in a bipolar asymmetric fashion with the bigger cluster at the lagging pole. However, the GAP activity of MglB is speculated to be dominating at the lagging pole. This is further stimulated by the presence of an MglB activator, RomY at the lagging pole^13^. It will be interesting to decipher the role of the MglB-Ct helix in MglA activation in presence of this co-GAP, RomY. Further, there is a possibility that the GAP and GEF domains of MglB are spatially regulated by interaction with other interactors such as MglC^25,26^. Recent reports show conformational changes in MglB dimers when complexed with MglA/RomY and MglC^27^. These could be critical in regulating the GAP activity of MglB at the lagging pole. Further, there could be yet unknown factors sequestering the GEF active MglB_1_ Ct-helix, spatially resolving its GAP and GEF functions. Disengaging the Ct-helix of MglB is essential to separate the GAP and GEF activities of MglB, thereby retaining MglB as a GAP only. In such a disengaged state, which could be driven by post-translational modifications or other interaction partners within the cell, MglB loses its GEF function and also detaches from MglA once GTP hydrolysis takes place. Together, this can facilitate the exclusion of MglA from the lagging pole and promote its recruitment to the leading pole and the Agl–Glt complexes by the RomR-RomX, MglB and other effectors.

Allostery is the mechanism of activation of an enzyme by interacting at a region other than its active site. In this case, through our design of linked complexes of MglAB, we conclusively prove that MglB is an allosteric activator of nucleotide exchange from MglA. Our study validates that the Ct-helix of MglB_1_ interacts with the helix binding pocket consisting of the β_2_-β_3_ loop of MglA which is the allosteric regulator site, thereby stabilizing the flipped conformation of MglA following the β-screw movement. Such a conformation of MglA possesses a nucleotide-binding pocket in an ‘open’ conformation ready to accommodate GTP, and the GDP dissociates easily. Consequently, when GTP binds, it associates tightly with the catalytic pocket which is further hydrolysed by the GAP activity of MglB. This is a unique mechanism - most of the GEFs of eukaryotic GTPases interact directly with the residues of the GTPase active site ^16^ to drive nucleotide exchange. It is possible that many mechanistically uncharacterized GEFs of eukaryotic small Ras-like GTPases could also act by an allosteric mechanism involving the inter-switch region. We predict that the families of small Ras-like GTPases which exhibit an inter-strand movement such as Arf, septins, and Rag GTPases are likely to exhibit these allosteric mechanisms of GEF activity^19,28,29^.

## Materials and Methods

### Cloning

Restriction-free cloning ^30^ was used to generate the constructs of MglA-link-MglB with C-terminal hexahistidine tag in *pHis17*-*Kan^R^* (*mglA-link-mglB-H_6_,* refer Addgene plasmid #78201 for vector backbone, *amp^R^* cassette replaced with kanamycin resistance using specific primers: pHis17 *Kan^r^*). Further MglA-link-MglB^ΔCt^ with C-terminal hexahistidine tag in *pHis17*-*Kan^R^* was cloned (*mglA-link-mglB^ΔCt^-H_6_*). The primers used are enlisted in Table 3. Similarly, MglB and MglB^ΔCt^ were cloned with a C-terminal Strep-tag in pHis17-*Amp^R^*. MglA and MglB constructs used were as reported in ^11^. The clones were subjected to *DpnI* (New England Biolabs Inc.) digestion followed by transformation. Positive clones were selected on a suitable antibiotic (kanamycin and ampicillin respectively) containing plates and checked by the consequent release of correct-sized fragments following double digestion with *NdeI* and *BamHI* (New England Biolabs Inc.). All the clones were confirmed by sequencing.

**Table 3:**
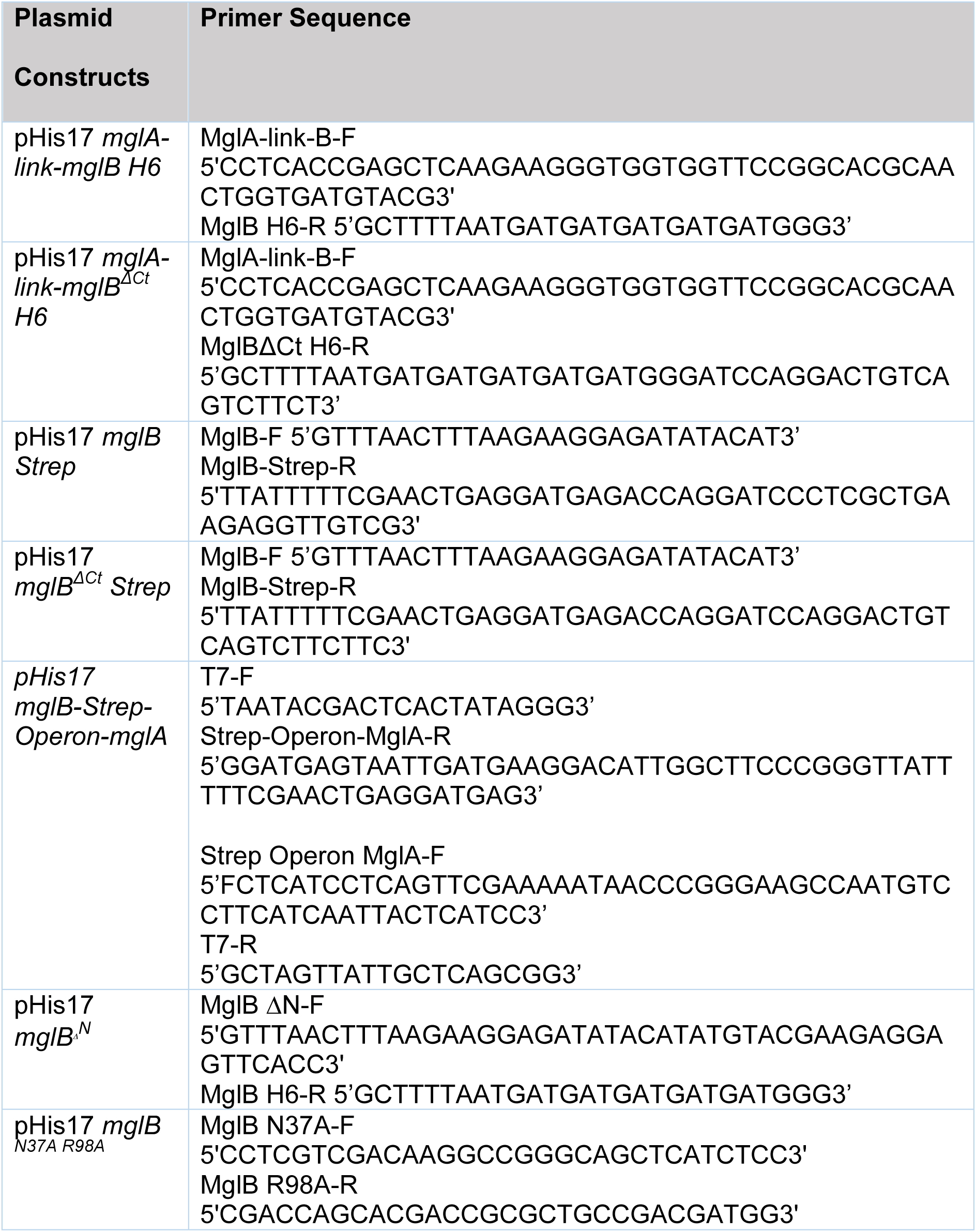
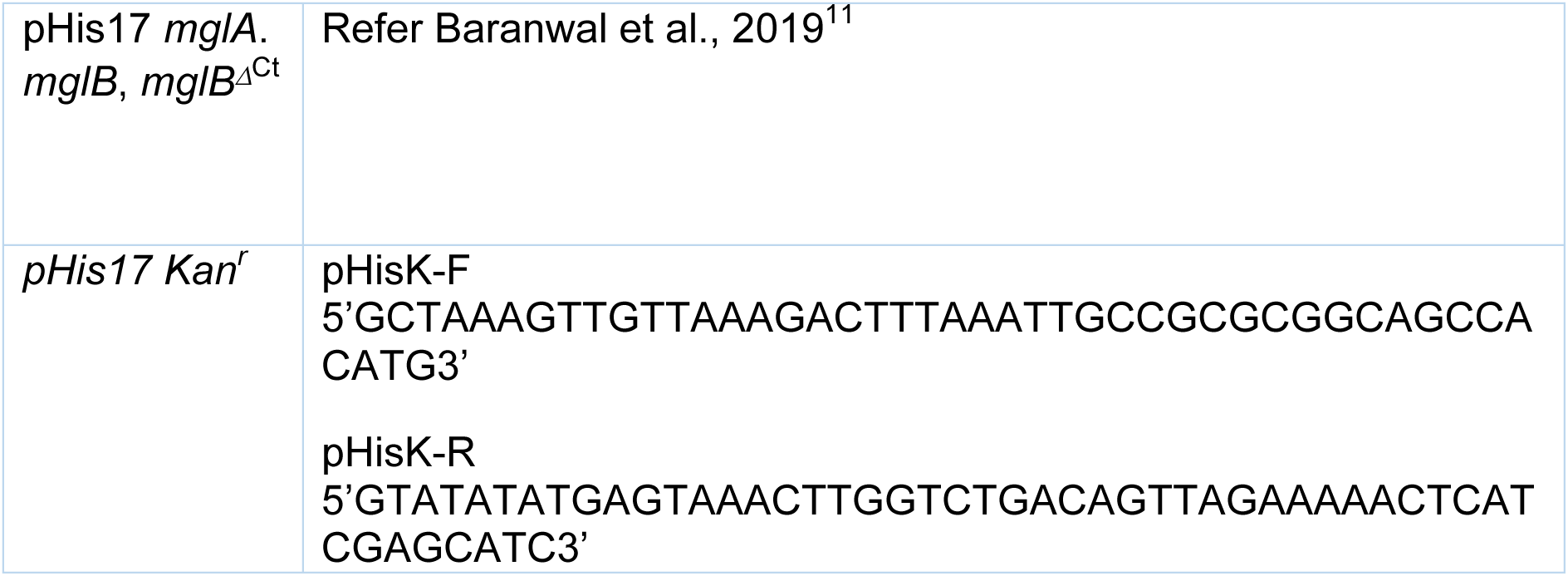
List of primers used

Firstly, MglB-Strep and MglA with the intervening operon sequence was cloned (*mglB-Strep-Operon-mglA)*. The operon sequence was synthesized using forward and reverse primers (Table 3) using overlap extension (Refer to Fig. S1 for details). This product was used as a megaprimer to be inserted in *pHis17 mglA-link-mglB-H_6_* construct as the template at the 5’ end of the *mglA* sequence. MglA has an internal *XhoI* restriction site. We performed double digestion using the internal *XhoI* and *HindIII* (on the vector backbone at the 3’ end of the gene) restriction sites on *mglB-Strep-Operon-mglA* and *mglA-link-mglB-H_6_* constructs to generate the vector and insert, respectively, for the ligation reaction (enzymes from New England Biolabs Inc.). *mglB-Strep-Operon-mglA* vector treated with TSAP (Promega) and the digested *mglA-link-mglB-H_6_* insert was ligated at the MglA *XhoI* restriction site using T4 DNA Ligase, (New England Biolabs Inc). The clones were transformed and screened similarly using double digestion with *NdeI* and *HindIII*. This generated the {AB^WT^} construct.

For {AB_1_^ΔCt^B_2_}, *mglB^ΔCt^-Strep-Operon-mglA* was PCR amplified using the same overlap extension primers as before, using *mglB^ΔCt^-Strep* construct. This product was digested using N-terminal *NdeI* and internal *XhoI* and inserted into a similarly digested *mglA-link-mglB-H_6_* vector. {AB_1_B_2_^ΔCt^}, and {AB^ΔCt^} were generated using combinations of *NdeI* and *XhoI* digested inserts of *mglB-Strep-Operon-mglA* and *mglB^ΔCt^-Strep-Operon-mglA*, respectively, and ligated to a digested *NdeI* and *XhoI mglA-link-mglB^ΔCt^-H_6_* template.

MglB^ΔN^, and MglB^N37,R98A^ mutations were incorporated in MglB using primers listed in Table 3. The clones were subjected to *DpnI* (New England Biolabs Inc.) digestion followed by transformation. Positive clones were selected on ampicillin-containing plates and checked by the consequent release of correct-sized fragments following double digestion with *NdeI* and *BamHI* (New England Biolabs Inc.). Both clones were confirmed by sequencing.

### Protein Expression

The plasmids were transformed in BL21DE3 (for {AB^WT^}, {AB_1_^ΔCt^B_2_}, {AB_1_B_2_^ΔCt^}) and BL21AI (for {AB^ΔCt^}, MglB^ΔN^, MglB^N37R98A^) strains of *E. coli* respectively. The cultures were grown in respective antibiotic-containing media (1X LB with 0.1 mg/ml ampicillin or 0.05 mg/ml kanamycin) and were kept in shaking conditions at 37°C. The cultures were induced with 0.5 mM IPTG (for BL21DE3) and 0.2% L-Arabinose (for BL21AI) once it reached the exponential phase of growth (between 0.6-0.8 OD). The cultures were incubated overnight at 18°C. The samples were subjected to 15% SDS-PAGE to observe the overexpressed band of protein of interest.

### Protein purification

Purifications of MglA and MglB were performed as described in reference 11^11^

#### His-tagged purification

For the MglAB linked constructs, harvested cells were resuspended in the lysis buffer L (50 mM Tris pH 8.0, 200 mM NaCl and 10% glycerol) at 4 °C. Consequently, the samples were centrifuged at 39,000 g for 45 minutes. The lysate was loaded on a 5 ml HisTrap HP (GE Healthcare) column. The loading buffer used was 200 mM NaCl, 50 mM Tris pH 8.0 and the elution buffer used was 200 mM NaCl, 50 mM Tris pH 8.0, 500 mM Imidazole. Protein was eluted using a stepwise gradient of 2%, 5%, 10%, 20%, 50%, and 100% of elution buffer.

#### StrepTrap

Elute from HisTrap consisting of the respective protein fractions were centrifuged at 39,000 g for 15 minutes and the supernatant was loaded on a 5 ml StrepTrap HP column (GE Healthcare). The binding buffer used was 150 mM NaCl, 50 mM Tris pH 8.0 and the elution buffer used was 150 mM NaCl, 50 mM Tris pH 8.0, 2.5 mM desthiobiotin. Elution buffer was injected into the column in 2 rounds of 5 ml each (Fraction I and II, respectively). The fractions with the protein were pooled, concentrated and injected into Superdex-75 column (GE Healthcare) to elute with a final buffer containing 50 mM NaCl, 50 mM Tris pH 8.0 (A50).

In the case of {AB^ΔCt^}, only HisTrap and StrepTrap were performed followed by washing off desthiobiotin using 50mM NaCl, 50mM Tris pH 8.0 (A50) buffer during protein concentration and consequently stored.

#### Size Exclusion chromatography

The column was equilibrated with 50mM NaCl, 50mM Tris pH 8.0 (A50) buffer. Concentrated protein after StrepTrap step (approximately 2-3 mg/ml, 200 µL of protein for analytical runs) was injected into the Superdex 75 size exclusion column (GE Healthcare) (volume less than 900 µL). UV absorbance at 280 nm was observed to monitor the elution of the protein. The respective fractions were pooled, concentrated and stored.

#### Anion Exchange (MonoQ)

For MglB mutants, the protocol was followed the same as MglB purification. However, following Superdex 75, the protein was impure. Hence it was injected in MonoQ 4.6/100 PE (GE Healthcare). Buffers used for binding and elution were Buffer A (50 mM Tris [pH 8.0], 50 mM NaCl) and Buffer B (50 mM Tris [pH 8.0], 1 M NaCl), respectively. A linear gradient of Buffer A ranging from 0% to 50% Buffer B over 20 column volumes was injected and the fractions containing the protein were pooled and concentrated.

### SEC-MALS

Size exclusion chromatography coupled with Multi-angle light scattering (SEC-MALS) experiments enabled the accurate mass estimation of the linked complexes. Superdex 200 Increase 10/300 GL column was used for SEC which was connected to an Agilent HPLC unit having an 18-angle light scattering detector (Wyatt Dawn HELIOS II) and a refractive index detector (Wyatt Optilab T-rEX). The experiments were performed at room temperature. The column was equilibrated with A50 (50 mM NaCl, 50 mM Tris pH 8.0) buffer at 0.4 ml/min. BSA at 2 mg/ml was used for the calibration of the system. The purified linked complexes (approximately 5 mg/ml, 110 µL) were consequently loaded to estimate the molecular weight of the eluted peaks. The Zimm model implemented in ASTRA software was used for the curve fitting and estimation of molecular weights. GraphPad Prism was used to average molar mass from fitted plots.

### NADH-coupled GTP hydrolysis assay

NADH coupled enzymatic assay ^31^, similar to the protocol used in Baranwal, et al, 2019^11^, was used to measure GTP hydrolysis activity. A master mix was prepared in A50 buffer (50 mM NaCl, Tris pH 8.0) containing 600 µM NADH, 1 mM phosphoenol pyruvate, 5 mM MgCl_2_, 1 mM GTP and pyruvate kinase and lactate dehydrogenase enzyme mix (∼25 U/ml). All components were mixed to a 200 µl reaction volume and added to Corning 96 well flat bottom plate. The reactions were initiated by adding purified complexes or a mix of MglA/MglB (mutants) in 1:2 ratios to a final concentration of 10 µM MglA. A decrease in NADH absorbance was measured at 340 nm using a Varioskan Flash (4.00.53) multimode plate reader. The absorbance was measured every 20 s for 7200 s. The initial time point and absorbance of buffer components were subtracted from all readings. NADH absorbance was converted to GDP produced using a slope obtained from a standard curve containing known concentrations of NADH. GraphPad Prism was used for data analysis and plotting the *k_obs_* values.

### Nucleotide exchange assay

The kinetic measurements for the linked complexes were performed on Fluoromax-4 (Horiba) where the intensity of fluorescence emission by *mant-*labelled GDP/GMPPNP (Jena Bioscience, Germany) at 440 nm was monitored after the excitation at 360 nm (protocol similar to the one used in Baranwal, et al, 2019^11^). The sample volume was 200 µl in a quartz cuvette (10 × 2 mm path length) and excitation and emission slit widths of 2 nm. *m*-GDP/GMPPNP (final concentration 800 nM) was present in buffer A50 (50 mM Tris pH 8.0, 50 mM NaCl, 5 mM MgCl_2_).

The protein mix, i.e., a final concentration of 3 µM of the respective linked complexes was added in the cuvette at 400 seconds after stabilization of the signal from only *mant*-nucleotide. Consequently, the fluorescence was recorded for 1,400 seconds where the increase in fluorescence intensity reflected the nucleotide-binding kinetics. At 1,800 seconds, the *mant-*labelled nucleotide was competed out with an excess of unlabelled GDP/GTP (final concentration 500 µM), resulting in the release of *mant*-labelled nucleotide from the protein which manifested as a decrease of fluorescence intensity. For plotting the relative intensities from the measurements, each value was divided by the average of the first 200 readings (400 seconds). These accumulations and decay reactions were fitted to exponential binding equations as given below in GraphPad Prism to estimate the *k_on_* and *k_off_* values.

P + N_1_ → PN_1_ + N_2_ → PN_2_ + N_1_, where P represents protein, N_1_ is the labelled nucleotide, N_2_ is the unlabelled nucleotide, and PN denotes the protein-nucleotide complex.

For estimation of *k_on_*, PN_t_ = PN_max_ (1 − e^−kt^).

For estimation of *k_off_*, PN_t_ = PN_max_ − PN_min_ (e^−kt^) + PN_min_

Here, PN_t_ represents the amount of the complex at time t, PN_max_ is the maximum amount of the complex upon association, and PN_min_ is the minimum amount of the complex after dissociation.

### High-Pressure Liquid Chromatography

The protein samples (100 µM each) were diluted with 50mM NaCl, 50mM Tris pH 8.0 buffer and then heat denatured at 75 °C. The heated protein sample was spun at 21,000g for 10 min. Then the supernatant was filtered with a 0.22-µm cellulose acetate filter (Corning), and the sample was loaded on the DNAPac PA200 ion exchange column (Thermo Fisher). Buffer A (2 mM Tris (pH 8.0)) was used as a binding buffer. The runs were performed at a flow rate of 0.4 ml/min, with linear gradients of 0% to 20% Buffer B (2 mM Tris (pH 8.0), 1 M NaCl) for 3 CV (column volumes), 20%–50% Buffer B for 3 CV and 100% B for 2 CV. The sample injection volume was 100 µl. The absorbance at 260 nm, indicative of the presence of GDP, was plotted against conductivity. The GDP standard was run with 100 µl of a 500 µM filtered GDP solution.

## Acknowledgements

The work was supported by the SERB Extramural Research grant (EMR/2014/000563), INSPIRE Faculty Research Grant (IFA12-LSBM-52) and common instrumentation facilities at IISER Pune. MK acknowledges INSPIRE and SC acknowledges IISER Pune, Infosys Foundation, Biochemical Society, UK and DBT (DBT/CTEP/02/20220753076) for fellowships. Dr Radha Chauhan and Sangeeta Niranjan from National Centre for Cell Science, Pune are acknowledged for their assistance with the use of the SEC-MALS facility. Suman Pal and Prof. Jayant Udgaonkar are acknowledged for their assistance with mass spectrometry.

## Author Contributions

SC: Designed and performed all the experiments and wrote the manuscript

MK: Performed cloning and initial purification of MglB mutants MglB^N37R98A^ and MglB^ΔN^ and reviewed the manuscript.

PG: Conceptualized the study, supervised the experiments and analyses, and reviewed and edited the manuscript

CrediT:

SC: Methodology; Investigation; writing - original draft;

MK: Investigation; Writing - review and editing

PG: Conceptualization; Methodology; Funding acquisition; Project administration; Supervision; Writing - review and editing

## Abbreviations

GAP: GTPase Activating Protein
mant: 2’/3’-O-(N-methyl anthraniloyl;
GDP: guanosine di-nucleotide;
GMPPNP: guanosine-5’-[(β,γ)−imido]triphosphate;
GTP: guanosine 5’-tri-phosphate;
GTPγS: guanosine:5’-O-[gamma-thio]triphosphate;
LDH: lactate dehydrogenase;
PCR: Polymerase Chain Reaction;
PDB: Protein Data Bank;
Pi: inorganic phosphate;
PK: pyruvate kinase;
MglA: Mutual gliding protein A;
MglB: Mutual gliding protein B,
NADH: nicotinamide adenine dinucleotide.

