## Supplementary Figures for "Mechanism of GTPase activation of a prokaryotic small Ras-like GTPase MglA by an asymmetrically interacting MglB dimer"

Indian Institute of Science Education and Research Pune, Dr Homi Bhabha Road, Pashan, Pune, India 411008.

This file includes Supplementary Figures S1 to S3

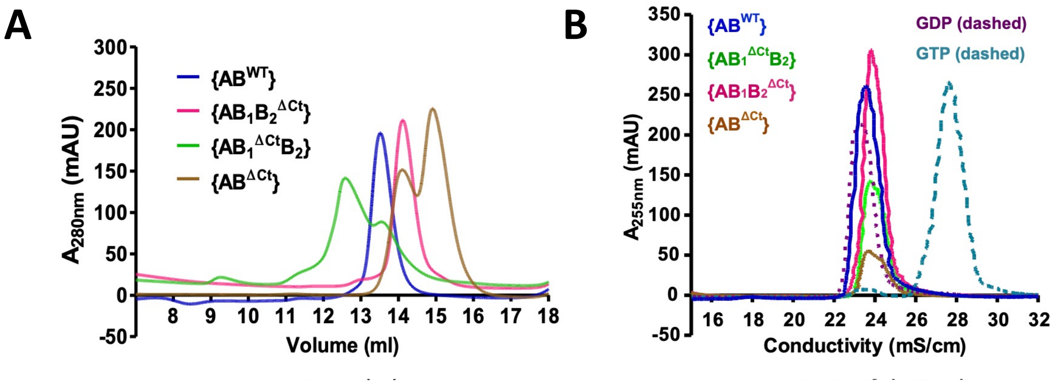


**Figure S1. SEC and HPLC profiles of linked complexes**

**A:** Analytical size exclusion chromatography (Superdex 200) profiles for the linked complexes: $\{AB^{WT}\}$ (blue), $\{AB_1B_2^{\Delta Ct}\}$ (magenta), $\{AB_1^{\Delta Ct}B_2\}$ (green) and $\{AB^{\Delta Ct}\}$ (orange).

**B:** HPLC profiles of the linked complexes: $\{AB^{WT}\}$ (blue), $\{AB_1B_2^{\Delta Ct}\}$ (magenta), $\{AB_1^{\Delta Ct}B_2\}$ (green) and $\{AB^{\Delta Ct}\}$ (orange).  GDP and GTP used as standards are shown in dotted lines in purple and cyan respectively.

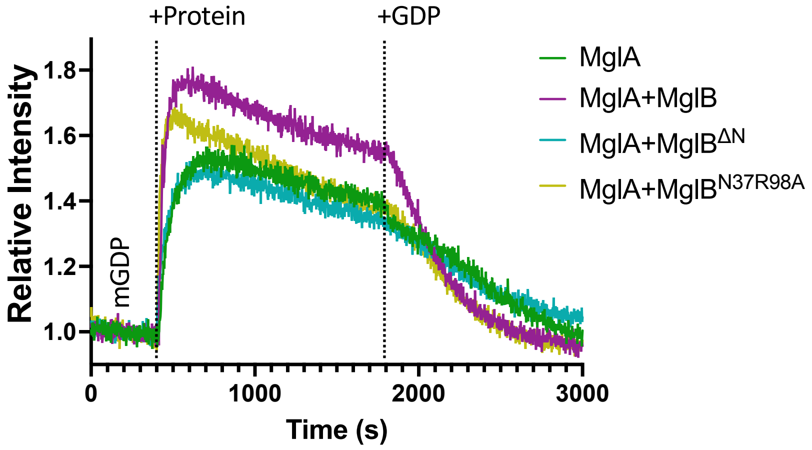


**Figure S2. Nucleotide exchange kinetics of MglA with excess of MglB mutants**

Kinetics of increase in *mant*-GDP fluorescence (region labelled as *m*-GMPPNP) upon adding MglA (dark green), MglA with MglB in 1:4 ratio (purple), MglB$^{\Delta N}$ (cyan) and MglB$^{N37R98A}$ (ochre) at 400 seconds (marked by dashed line labelled '+P'; Stage I), followed by competition of *mant*-GDP by adding excess unlabelled GDP at 1800 seconds (marked by dashed line labelled '+GDP'; Stage II).

The phases I and II, which represent association and dissociation for $k_{on}$ and $k_{off}$ estimation respectively (tabulated in Table 1B), are demarcated by solid line.

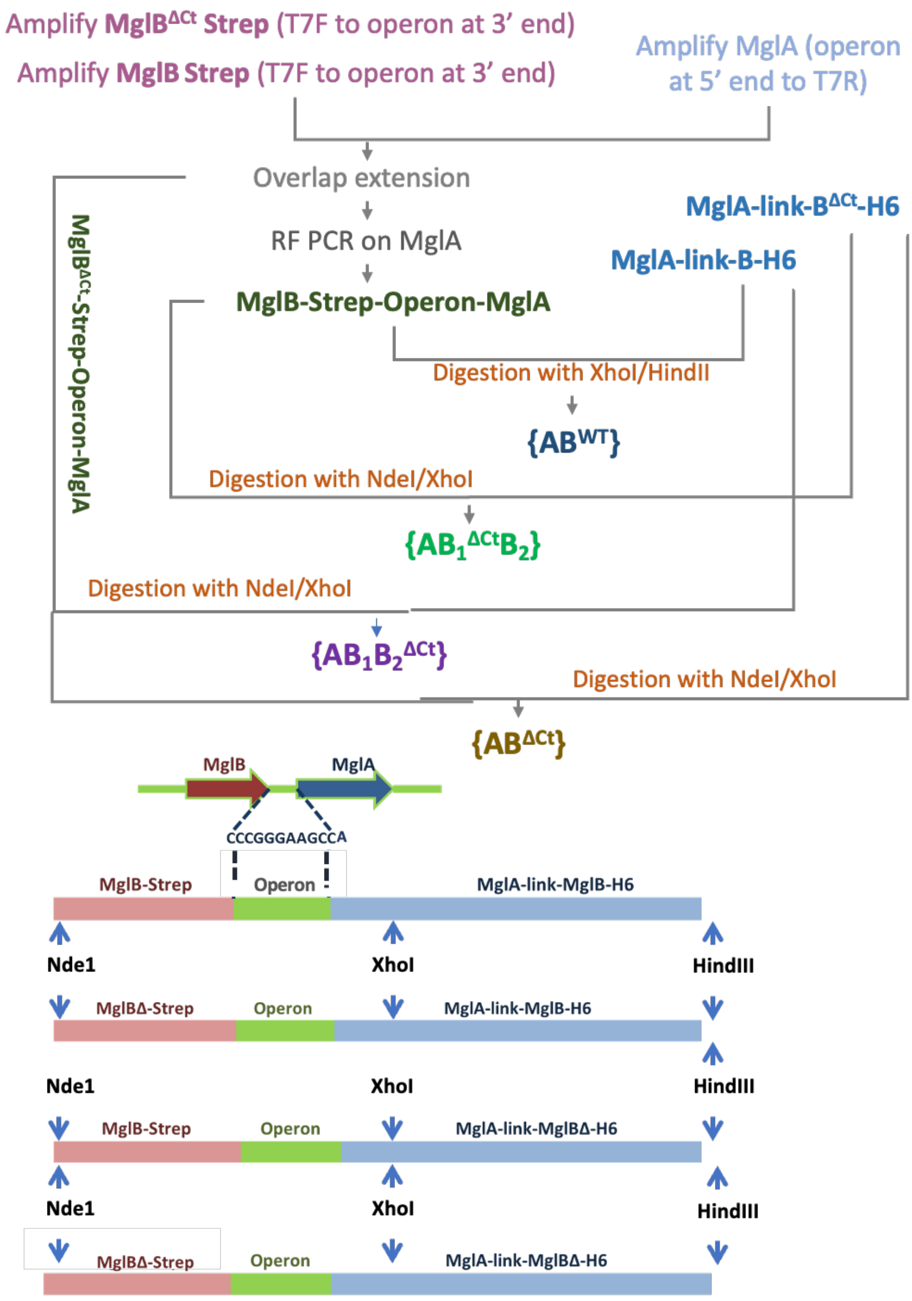


**Figure S3. Strategy for cloning the asymmetric linked constructs of MglA and MglB**

*mglB* and *mglA* genes are present in the same operon in the *Myxococcus xanthus* genome with an intervening sequence. These two loci were replaced with MglB-Strep (forming the MglB$_2$ protomer) and the MglA-link-MglB$_1$, respectively. The steps for cloning are detailed with the relevant restriction sites highlighted. Refer to Table 1 for the respective primers and construct details.
